# CDK12 controls transcription at damaged genes and prevents MYC-induced transcription-replication conflicts

**DOI:** 10.1101/2024.06.03.597073

**Authors:** Laura Curti, Sara Rohban, Nicola Bianchi, Ottavio Croci, Adrian Andronache, Sara Barozzi, Michela Mattioli, Fernanda Ricci, Elena Pastori, Silvia Sberna, Simone Bellotti, Anna Accialini, Roberto Ballarino, Nicola Crosetto, Mark Wade, Dario Parazzoli, Stefano Campaner

## Abstract

Oncogene-induced replicative stress is a potent tumor-suppressive mechanism that must be kept in check for cancer cells to thrive. Thus, the identification of genes and pathways involved in replicative stress is key to understand cancer evolution and to identify prospective therapeutic targets. Here, we investigated factors that modulate replicative stress upon deregulation of the MYC oncogene. We identified the cyclin-dependent kinase CDK12 as selectively required to prevent transcription-replication conflicts and the activation of a cytotoxic DNA-damage response (DDR). At the mechanistic level, CDK12 was recruited to damaged genes by PARP-dependent DDR-signaling and elongation-competent RNAPII. Once recruited, CDK12 repressed transcription by preventing the association of CDK9 with RNAPII. Either loss or chemical inhibition of CDK12 led to DDR-resistant transcription at damaged genes. Genome-wide profiling revealed that loss of CDK12 exacerbated transcription-replication conflicts in MYC-overexpressing cells and led to the accumulation of double-strand DNA breaks (DSBs), occurring preferentially between early-replicating regions and transcribed genes, organized in a co-directional head-to-tail orientation. Overall, our data demonstrate that CDK12 protects genome integrity by repressing transcription of damaged genes, which is required for proper resolution of DSBs at oncogene-induced transcription-replication conflicts. This provides a rationale that explains both how CDK12 deficiency can promote tandem duplications of early-replicated regions during tumor evolution, and how CDK12 targeting can exacerbate replicative-stress in tumors.

## INTRODUCTION

Replication stress (RS) is a major hallmark of cancer cells and a source of genomic instability^1^. During tumor progression, mutations in oncogenes and tumor suppressor genes can induce RS by imposing a state of high and deregulated DNA replication, which increases the chances for fork collapse and generation of DNA breaks^2^. In this scenario, fork collapse has been linked to multiple causes, including the depletion of nucleotides, mis-incorporation of ribonucleotides into nascent DNA, inefficient activation of protective fork processing pathways and conflicts arising from the encounter of replication and transcription complexes^3–5^.

RS is considered to have a dual role in cancer evolution: on one hand, it can act as a driving factor by providing a mutation-prone background that favors further selection of driver mutations; on the other, it can trigger tumor suppressive responses such as apoptosis or cellular senescence^6^. In line with the latter, the exacerbation of RS in cancer cells can offer therapeutic opportunities, either in combinatorial treatments, to enhance effectiveness of current chemotherapeutic drugs, or in targeted therapies exploiting synthetic-lethal dependencies in genetically defined tumors^7–9^. Thus, identifying factors and processes that regulate RS in cancer cells is a subject of active studies and has high translational potential.

The c-MYC proto-oncogene is frequently overexpressed in cancer cells, due to genomic alterations (e.g., translocation, amplification) or oncogenic mutations in upstream regulatory pathways^10,11^. Its product, MYC, is a transcription factor belonging to the bHLH-LZ family. It dimerizes with another bHLH-LZ protein, MAX, to bind DNA in a sequence specific manner and to control the transcription of genes linked to cell proliferation and metabolism^11^. Oncogenic levels of MYC lead to increased binding to proximal and distal regulatory elements within the genome, a phenomenon dubbed “enhancer-promoter invasion”, that pairs with broadened transcriptional regulation^12–15^. The essentiality of MYC in cancer cells and its temporal dispensability in normal tissues^16^, makes it an attractive and potentially universal therapeutic target^17^, whether through direct inhibition of MYC activity^18^, or through exploitation of MYC-induced dependences^18^.

While still under investigation, evidence shows that MYC can trigger RS by promoting processivity of DNA synthesis, increasing origin firing, anticipating S-phase entry and possibly by inducing transcription-replication conflicts (TRCs)^19–21^. Several lines of evidence indicate that MYC-overexpressing cells are addicted to genetic dependencies that curb RS to avoid rampant genomic instability and cell death. These include the ATR/CHK1 kinases^22–25^, fork remodelers and regulators^26–28^, and factors preventing transcriptional interference^25,29–33^. Altogether, these pathways may cooperate to prevent collapse and/or excessive processing of stalled replication forks.

To identify factors regulating genomic and genetic stability in MYC overexpressing cells we conducted a loss of function genetic screen. Among others genes, we identified CDK12 as being selectively required to prevent RS and cytotoxic DDR. CDK12 is a member of the cyclin dependent kinase family that associates with and is activated by Cyclin K (CCNK)^34^. Like other transcriptional CDKs, CDK12 - and its paralogue CDK13 - can phosphorylate and activate RNA polymerase II (RNAPII)^35^. In addition, CDK12 regulates the expression and processing of genes linked to cell cycle progression and DNA damage signaling and repair^36–38^. In line with this, inhibition of CDK12 sensitizes cancer cells to DNA-damaging agents^34^. CDK12 is also recurrently mutated in several cancers, and an actionable therapeutic target^34,39^.

Here, we report that CDK12 is recruited onto transcribed and damaged genes to repress transcription. This depends on PARP, engaged by DDR signaling, and the elongation factor SPT5. Inhibition or silencing of CDK12 unleashes transcription at damaged genes and exacerbates TRCs leading to DSBs between early replicating regions and the promoters of genes transcribed co-directionally to the replication fork. Our work unveils a new role for CDK12 in controlling transcription and genomic stability and reveals how management of TRCs is a major liability in MYC-driven tumors.

## RESULTS

### Identification of genes controlling genome stability and viability in MYC-overexpressing cells

To identify modulators of genomic stability and cell viability in MYC-overexpressing cells, we devised a high-content siRNA screen in Rosa26-MycER mouse embryo fibroblasts, where conditional activation of MycER mimics the oncogenic properties of c-MYC^27^. These cells were generated from homozygous knock-in embryos carrying a gene encoding a MYC-estrogen receptor chimera (MycER) that can be activated by 4-hydroxytamoxifen (OHT)^40^. We screened this cell line with a custom siRNA library (DDR-library) targeting 1196 genes previously implicated in genome integrity^41^, and a commercial library (druggable-library) targeting 1400 druggable genes. For the screen, cells were retro-transfected in 384-well plates, with each well containing a siRNA mix targeting one gene. Forty-eight hours post-transfection, cells were analyzed by immunofluorescence for a parallel quantification of DDR signaling (based on the integration of single-cell DH2AX intensities) and cell growth (by cell counting), both in mock (no OHT) and

### MycER-activated cells (plus OHT)

For each gene, we calculated the average normalized differential values (Z-score) for both DDR and cell growth (Fig. 1a, Extended Data Fig. 1a,b,d,e and Supplementary Table 1) and identified as hits those genes that, when silenced, would preferentially affect DDR (DDR hits) and/or cell growth (viability hits). We scored a total of 629 genes: 386 DDR-hits and 335 viability-hits; of these 92 were both DDR and viability hits (Fig. 1b and Extended Data Fig. 1c,f). A subset of the primary hits was cross validated in a secondary screen (Extended Data Fig. 1b,c,f and Supplementary Table 1). Gene ontology and protein-protein interaction network analyses with Metascape^42^, revealed a prevalence of genes involved in RNA processing, translation, protein degradation, cell cycle control and signaling (Fig. 1c, Extended Data Fig.2 and Supplementary Table 2). This confirmed previously identified MYC-dependencies, such as its reliance on transcriptional regulators^43^, the splicing machinery^44,45^, regulators of mitosis^46,47^, nucleotide biosynthesis ^48,49^, and proteasome activity ^50^.

**Figure 1.**
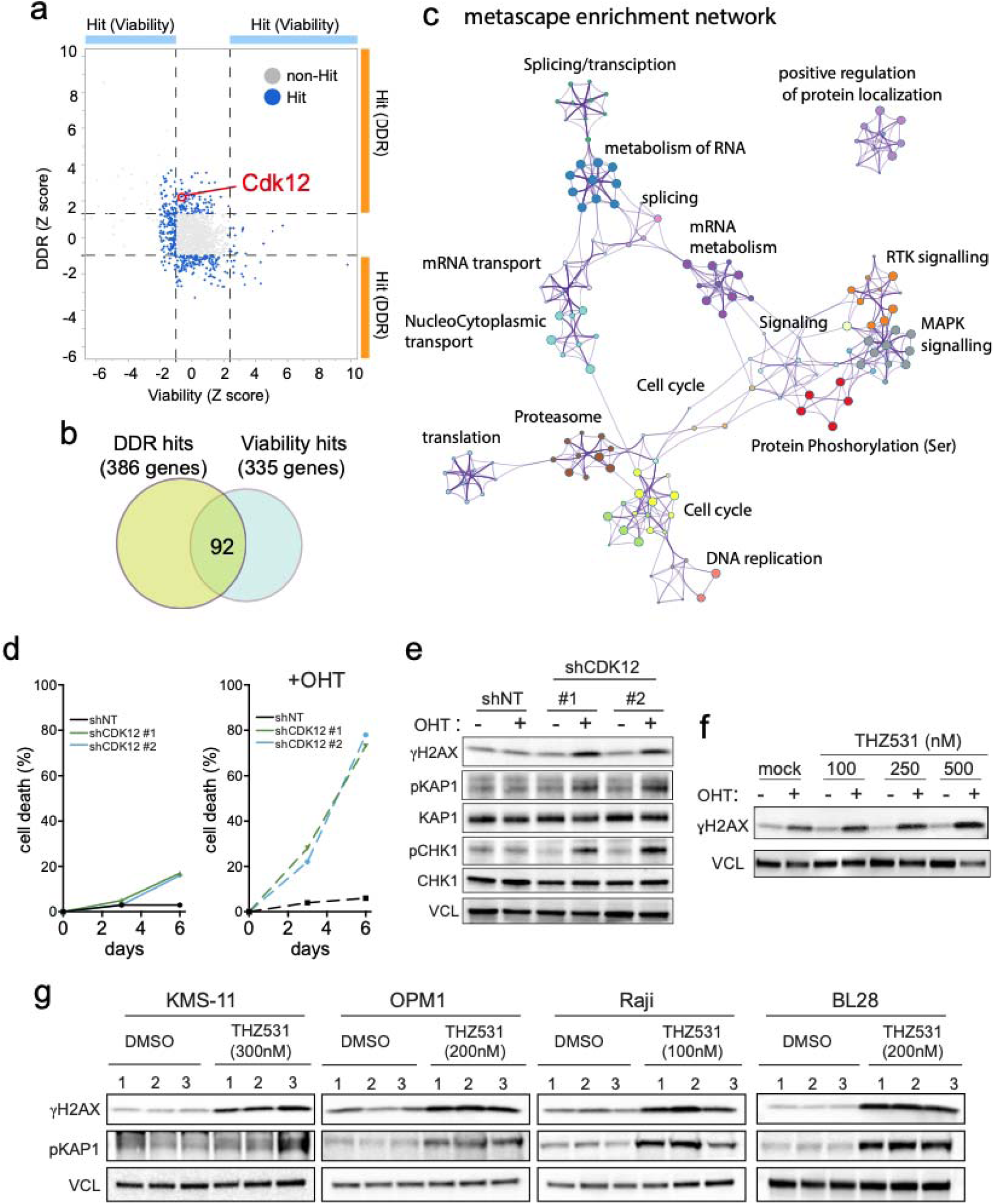
CDK12 is synthetic lethal with Myc activation. **a**, Dot plot of the normalized differential DDR score (y axis, Z-score for DDR) and the normalized differential viability (x axis, Z-score for viability) for each siRNA of the library targeting druggable genes (primary screen). A positive Z-score for DDR indicates higher DDR upon MycER activation compared to mock activated U2OS-MycER cells, while a negative viability Z-score indicates a reduction of viability compared to mock activated cells. Dashed lines indicate thresholds used to call the DDR and viability hits. **b**, Venn diagram of all the hits identified in the two primary screens as DDR- and viability-hits. **c**, Metascape enrichment network of the hits of the two primary screens (DDR and druggable genes). **d**, Cell death in U2OS-MycER cells upon MycER activation (+OHT, 400 nM) and CDK12 knockdown by the indicated shRNAs. **e,f,** WB analysis of DDR markers in U2OS-MycER cells upon MycER activation (+OHT, 400 nM) and (**e**) CDK12 knockdown or (**f**) CDK12 inhibition by THZ531. **g**, WB analysis of DDR markers following treatment of multiple myeloma (KMS-11, OPM1) and Burkitt’s lymphoma (Raji, BL28) cell lines with THZ531 for 48 hours. Vinculin (VCL) was used as loading control. Lanes 1, 2, and 3 show independent replicates.

Among the hits confirmed in the secondary screen (Extended Data Fig.1b), we identified CDK12, a kinase implicated in transcriptional control and genome stability, which has recently drawn attention as a potential therapeutic target in cancer^34,39^. Most noteworthy, CDK12 also scored in a previous list for MYC-synthetic lethal candidates, but without further characterization^47^. In addition, re-analysis of a recently published screen^51^ revealed that CDK12, its paralogue CDK13, and its regulatory CCNK are synthetic lethal with MYC amplification in medulloblastoma (Extended Data Fig. 3).

### Genetic or chemical targeting of CDK12 impairs cell viability and triggers a DNA damage response in MYC-overexpressing cells

For further validation and mechanistic analysis, we engineered U2OS cells expressing the MycER chimera^27^ (U2OS-MycER cells) with doxycycline-inducible shRNAs targeting CDK12 (shCDK12 #1 and #2) or a non-targeting shRNA (shNT) (Extended Data Fig.4a). CDK12 knock-down (CDK12- KD) impaired cell growth, with a stronger inhibition when MycER was activated (Extended Data Fig. 4b,c). On the other hand, cell viability was reduced only when cells experienced both CDK12- silencing and MycER activation (Fig. 1d), indicating a strong synthetic-lethal phenotype. This was paralleled by enhanced DNA damage response (DDR), engaging both ATM and ATR signaling, as indicated by WB analysis (Fig. 1e). Synthetic lethality and synthetic DDR activation were also confirmed by siRNA mediated silencing (Extended Data Fig. 5) and upon CDK12 inhibition by THZ531 or dinaciclib (Fig. 1f and Extended Data Fig. 6). Similar to CDK12, silencing of CDK13 or CCNK lead to synthetic lethality and DDR upon MycER activation (Extended Data Fig. 7). Next, we evaluated whether CDK12 inhibition would induce DDR in Burkitt’s lymphoma and multiple myeloma cell lines in which expression of MYC is deregulated by chromosomal translocations. Most of the cell lines tested were sensitive to THZ531, with IC50 values in the nanomolar range (Extended Data Fig.8a). Similar to what was observed in U2OS-MycER cells, THZ531 induced the activation of a DDR characterized by a recurrent increase of phospho-KAP1 and DH2AX signals (Fig. 1g). The only exception was the RAMOS cell line, which had the highest IC50 for THZ531 and did not display an increase in phospho-KAP1 or DH2AX following CDK12 inhibition, but instead showed increased phosphorylation of CHK1 (Extended Data Fig. 9). Other DDR markers had inconsistent fluctuations when these cell lines were treated with THZ531 (Extended Data Fig. 9), possibly suggesting that the engagement of select DDR pathways may depend on the genetic background or other context-specific properties of the cancer cells. In all the cell lines tested CDK12 inhibition also led to an altered cell cycle distribution characterized by a decrease in BrdU-labelled cells indicating reduced DNA synthesis and altered cell cycle progression (Extended Data Fig. 8b).

### Genome-wide transcriptional alterations upon CDK12 silencing

CDK12 has been implicated in transcriptional regulation as it can phosphorylate RNAPII on serine 2 of the carboxyl-terminal repeat domain (CTD), thus stimulating elongation^35^. It was also reported to prevent early transcriptional termination at cryptic poly-A sites and to regulate alternative splicing^37,38,52^. Thus, we assessed how CDK12 silencing would affect the transcriptome in our cells. RNA-seq analysis showed that CDK12-KD led to both up- and down-regulation of distinct sets of genes (DEG-up and -down, Extended Data Fig. 10a). These alterations in mRNA expression were coherent with phospho-Ser-2 RNAPII ChIP-seq data, with consistent increases and decreases in the elongating form of RNAPII on DEG-up and DEG-down genes, respectively (Extended Data Fig. 10b), while the global level of Ser2-Pi RNAPII was not affected by CDK12 silencing (Extended Data Fig. 10c), confirming previous observations^35,37,38^. Most noticeably, DEG-down genes tended to be more expressed in unchallenged cells and had a shorter size, while the opposite was true for DEG-up genes (Extended Data Fig. 10d,e and Supplementary Table 3). We also evaluated whether CDK12 silencing would broadly alter the expression of DDR genes. The top 20 GO enrichment terms of DEG did not include terms associated with genome stability or DNA repair (Extended Data Fig. 10f,g and Supplementary Table 3). The analysis of a custom collection of DDR associated genes did not reveal global repression of these genes (Extended Data Fig. 11a-c and Supplementary Table 3). Exon level analysis of the mRNA processing of long-DDR genes failed to show any consistent or overt alterations in splicing or premature termination (Extended Data Fig. 11d and 12). Overall, the lack of pervasive regulation of DDR genes following CDK12 silencing suggests that the increased DDR observed upon MYC activation in CDK12-KD cells may not be solely ascribed to altered expression or processing of some DDR genes. In addition, CDK12-KD did not alter MYC-dependent transcription, since MYC targets were still efficiently activated when CDK12 was silenced (Extended Data Fig. 13 and Supplementary Table 3).

### CDK12 represses RNA synthesis at damaged genes

Given the reported role of CDK12 in regulating RNAPII activity, we asked whether silencing of CDK12 might induce a global change in RNA synthesis rate, not necessarily detected by steady state expression analyses (RNA-seq). To this end, we pulsed labelled cells and evaluated RNA synthesis by quantitative immunofluorescence (IF) analysis of 5-ethynyluridine (EU) incorporation. We assessed RNA synthesis in unchallenged cells or following DNA damaging radiations (UV or IR), which are known to repress transcription. Silencing of CDK12 led to an increase in EU incorporation suggesting a raise in global RNA synthesis or, alternatively, an imbalance between genes repressed and genes activated by CDK12 (i.e. DEG-up genes would contribute more to global RNA synthesis than DEG-down). Surprisingly, while IR or UV irradiation caused a global decrease in RNA synthesis, the increase caused by CDK12-KD persisted in irradiated cells, as shown in multiple cell lines (Fig. 2a,b and Extended Data Fig. 14a,c). Inhibition of CDK12 by THZ531 yielded similar results (Extended Data Fig.14b). As increased RNA synthesis caused by CDK12 depletion persisted in IR- or UV-irradiated cells, we hypothesized that damaged genes may not be repressed in these conditions. A role for CDK12 in the repression of transcription at damaged loci would have important implication, as this DNA-damage induced transcriptional shutdown is critical for preserving genomic integrity^53^. To further address this, we took advantage of the U2OS-TRE-I-SceI-19 reporter cell line, that allows to introduce DNA double-strand breaks (DSBs) upstream of the promoter of a Tet-regulated MS2-based transcriptional reporter^54^. This system enables simultaneous locus-specific detection of nascent RNA and DSBs, at the single-cell level. Upon activation of the mCherry-tTA-ER transcription factor, the transcribed locus can be visualized by the colocalization of mCherry-tTA-ER and the nascent RNA, which is bound by the MS2-YFP RNA binding protein. At the same time, DSBs generated by the expression of the I-SceI restriction enzyme can be visualized by DH2AX IF-staining. As expected, upon expression of I-SceI, DSBs were generated at the locus (visualized by the colocalization of DH2AX foci with mCherry-tTA-ER) (Fig. 2d), thus leading to loss of nascent RNA (i.e. loss of the nuclear MS2-YFP dot, Fig. 2d). Indeed, less than 10% of mock silenced cells (siLuc) showed an MS2-YFP signal colocalized with DH2AX (Fig. 2e), thus confirming that upon DNA damage the activity of the transcriptional reporter was strongly suppressed. Instead, upon CDK12 silencing, transcription was rescued in 50% of the cells showing a damaged site (Fig. 2d,e). A similar rescue was observed upon inhibition of CDK12 by THZ531 (Fig. 2f,g). In both cases, loss of CDK12 activity does not affect the efficiency of DSBs induced by I-SceI (Extended Data Fig.15a,b). In the absence of I-SceI, mCherry-tTA-ER and MS2-YFP colocalized and were not affected by CDK12 silencing or inhibition (Fig. 2d,e,f,g). Altogether, these data imply that CDK12 activity is required for locus-specific transcriptional repression following DNA damage. Silencing of either CCNK or CDK13, also led to a partial rescue of transcription at the damaged TRE-I-SceI-19 reporter, while simultaneous silencing of CDK12, CDK13 and CCNK further increased transcriptional rescue of the locus, suggesting partial redundancy (Extended Data Fig. 16a,b).

**Figure 2.**
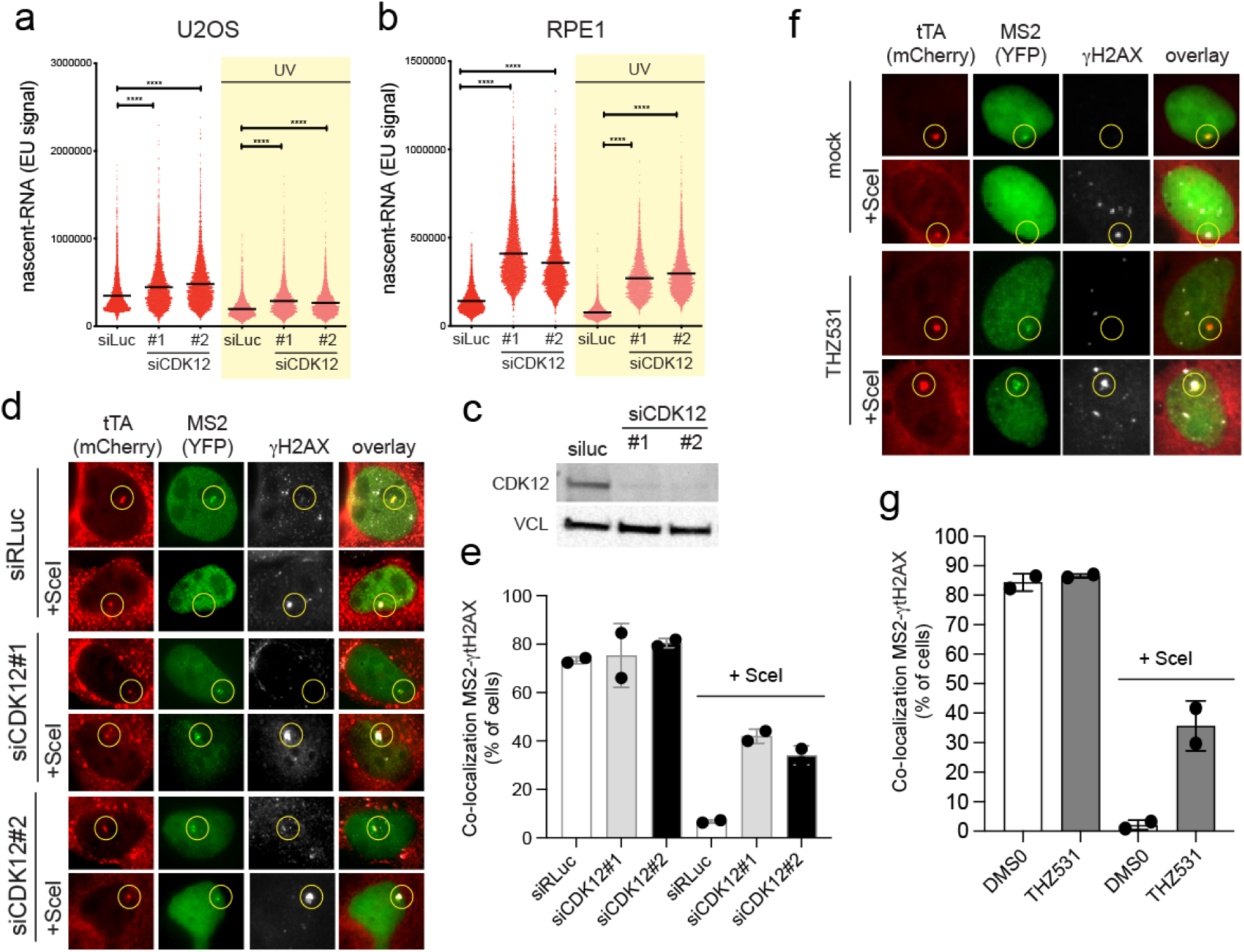
CDK12 silencing or inhibition rescues transcription at DNA damaged genes. **a**,**b**, Bee swarm plots of single-cell nascent-RNA synthesis (by EU incorporation) in mock or UV-irradiated cells. Cells were pulsed with 0.5 M EU for 20 min. before collection. (**a**) U2OS cells: siLuc n=1965; siCDK12#1: n=2320; siCDK12#2: n=3446; siLc+UV: n=2363; siCDK12#1+UV: n=5003; siCDK12#2+UV: n=2806. (**b**) RPE1 cells: siLuc n=4100; siCDK12#1 n=4090; siCDK12#2: n=4482; siLc+UV: n=5503; siCDK12#1+UV: n=2897; siCDK12#2+UV: n=4057. **c**, WB analysis. VCL is a loading control **d**,**e**, Representative IF-images (**d**) and bar plot (**e**) of the colocalization of mCherry-tTA-ER and YFP-MS2 signals upon silencing of CDK12 in U2OS-TRE-I-SceI-19 cells. Where indicated, cells were transfected with I-SceI (+SceI) to induce DSBs on the TRE-MS2 reporter. Average of two independent experiments. siRLuc: n=58, n=109; siCDK12#1: n=97, n=117; siCDK12#2: n=48, n=94; siRLuc+SceI: n=64, n=109; siCDK12#1+SceI: n=70, n=102; siCDK12#2+SceI: n=67, n=76. **f**,**g**, as in d,e, but upon inhibition of CDK12. Average of two independent experiments. DMSO: n=159, n=124; THZ531: n=123, n=93; DMSO+SceI: n=113, n=87; THZ531+SceI: n=111, n=96.

### CDK12 is recruited to damaged and transcribed genes

Next, we asked whether CDK12 could localize to sites of DNA damage. Cells were irradiated and proximity ligation assay (PLA) was used to assess the proximity of CDK12 to DH2AX foci. While signals were negligible in unirradiated cells, there was a marked increase in PLA-signals (foci) 30 minutes post-irradiation (Fig. 3a,b); by 4 hours the signal had returned to baseline level, possibly coinciding with ongoing DNA repair (Fig. 3a,b). The same result was confirmed with a different CDK12 antibody (Extended Data Fig. 17). Silencing of CDK12 lowered the PLA signal in irradiated cells, confirming the selectivity of the assay (Fig. 3a,b and Extended Data Fig. 17). Hence, CDK12 can be detected in close proximity to DDR-foci. To assess whether DNA damage and/or the resulting DDR would trigger the recruitment of CDK12, we transiently expressed an mCherry-tagged CDK12 and performed live imaging analyses on laser micro-irradiated cells. In unchallenged cells, CDK12 showed a diffused/dotted nuclear signal, while upon irradiation CDK12 promptly localized at damaged sites (Fig. 3c, Extended Data Fig.18a,b and Supplementary Video 1). Similarly, mCherry-CDK12 was recruited onto the reporter locus only when this was cut by the I-SceI nuclease (Fig. 3d and Extended Data Fig. 19a).

**Figure 3.**
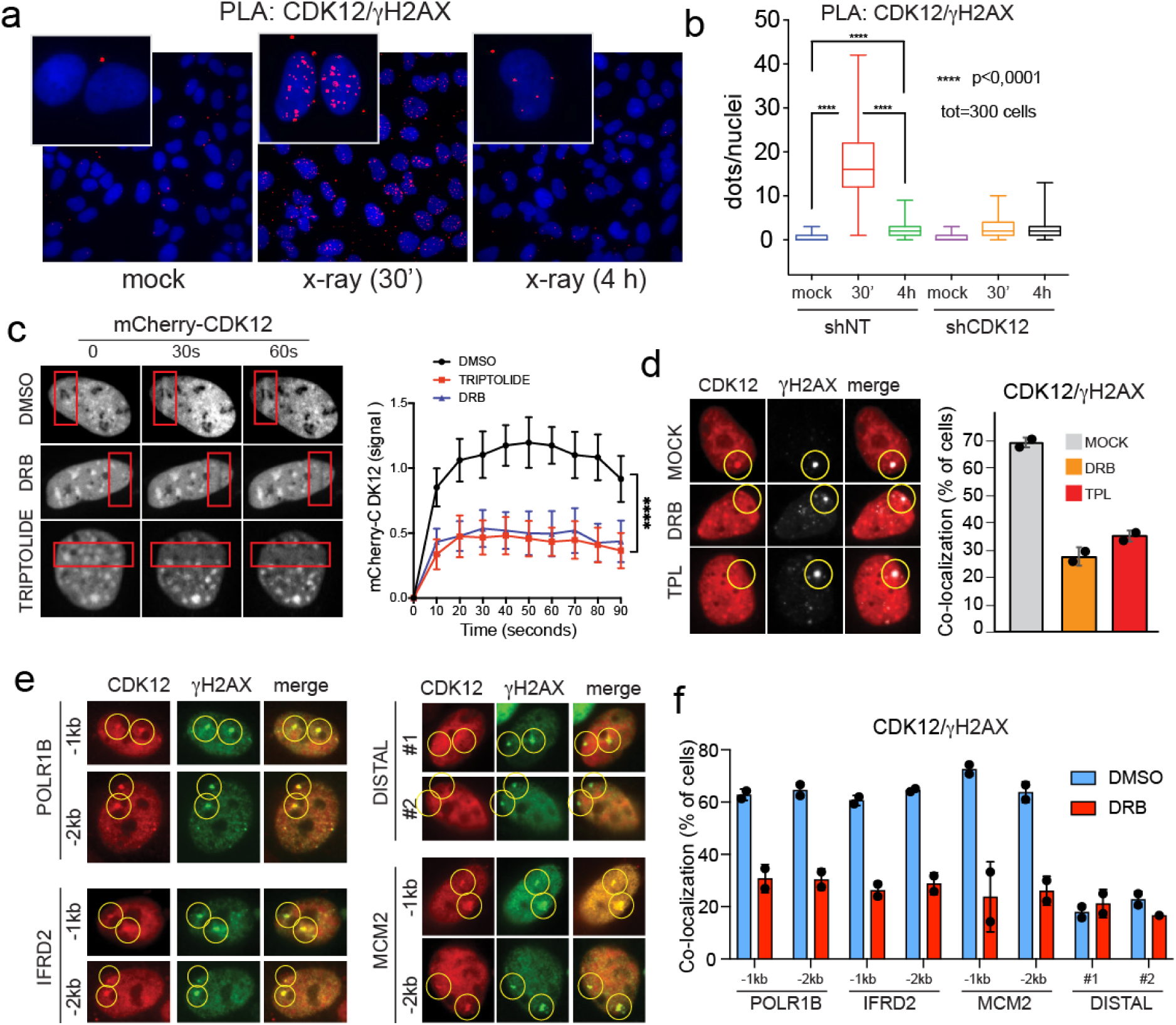
CDK12 is recruited on DNA damaged sites proximal to transcribed genes. **a**,**b**, Analysis of proximity of CDK12 and DH2AX foci by PLA in U2OS cells upon CDK12 silencing (shCDK12) or mock (shNT, non-targeting). (**a**) Representative pictures of mock silenced cells (shNT). shNT: mock n=284; 30’ n=317; 4h n=301. shCDK12: mock n=283; 30’ n=303; 4h n=310. (**b**) Box plot of PLA signals (nuclear dots) at different times post-irradiation (10 Gy). **c**, Kinetics of recruitment of mCherry-CDK12 to laser-damaged sites in U2OS cells treated or not with 100 µM DRB or 1 µM triptolide (TPL). Left, representative pictures; the red boxes highlight the irradiated areas. Right, time-series plot. DMSO: n=5; DRB: n=13; TPL: n=12. **d**, Colocalization of mCherry-CDK12 and DH2AX foci at SceI-induced DSBs in U2OS-TRE-I-SceI- 19 cells treated or not with 100 µM DRB or 1 µM triptolide (TPL). Left, pictures of mCherry-CDK12 and DH2AX signals. Right, box plots of two independent experiments. MOCK n=43, n=60; DRB n=52, n=61; TPL n=50, n=57. **e**,**f**, IF analysis of locus specific double strand breaks induced by CRISPR/Cas9 editing at select loci. (**e**) Representative images. (**f**) Bar plot reporting the average colocalization of CDK12 and DH2AX foci in DRB treated cells (average of two experiments). DMSO POLR1B -1kb: n=28, n=31; DRB POLR1B -1kb: n=37, n=26; DMSO POLR1B -2kb: n=23, n=32; DRB POLR1B: -2kb n=29, n=30; DMSO IFRD2: -1kb n=27, n=29; DRB IFRD2 -1kb: n=28, n=25; DMSO IFRD2 -2kb: n=23, n=25; DRB IFRD2 -2kb: n=12, n=27; DMSO MCM2 -1kb: n=24, n=35; DRB MCM2 -1kb: n=21, n=28; DMSO MCM2 -2kb: n=23, n=27; DRB MCM2 -2kb: n=18, n=30; DMSO DISTAL#1: n=19, n=30; DRB DISTAL#1: n=23, n=24.

Further analysis with chemical inhibitors revealed two important features of the CDK12 relocalization. First, CDK12 inhibition by THZ531 did not prevent its recruitment to laser-damaged sites, thus indicating that this was independent from CDK12 catalytic activity (Extended Data Fig.18c), yet this required CCNK (Extended Data Fig.16c). Second, the recruitment of CDK12 onto damaged sites in both microirradiated and I-SceI transfected cells was suppressed by the transcriptional inhibitors 5,6-Dichloro-1-beta-D-ribofuranosylbenzimidazole (DRB) or triptolide, indicating that it depended on RNAPII transcriptional activity (Fig. 3c,d). Similarly, lack of activation of rtTA and the consequent recruitment of the transcriptional machinery prevented the localization of CDK12 on the damaged reporter in I-SceI transfected cells (Extended data figure 19b). To verify CDK12 recruitment on endogenous DNA damaged loci, we took advantage of the CRISPR/Cas9 genome editing system and designed sgRNAs to generate DSBs either at the promoter of expressed genes (MCM2, IFDR2 and POLR1B) or at non-transcribed distal regions. While sgRNA- and Cas9-mediated induction of DSBs proximal to promoters induced two DH2AX foci that co- localized with CDK12, the latter was absent at DSBs induced at the non-transcribed distal sites (Fig. 3e,f). At transcribed loci, CDK12 localization depended on the activity of RNAPII, since it was reduced to background level by DRB (Fig. 3f). Also, CDK12 recruitment was low and near background levels when DSBs were induced by sgRNAs targeting the transcriptional termination sites of transcribed genes (Extended Data Fig.19c). Overall, these results suggest that CDK12 is recruited to DSBs that occur proximal to the promoter of transcribed loci or at gene bodies.

### PARP, but not ATM- or ATR-dependent DDR, controls the recruitment of CDK12 to damaged genes

Next, we asked which components of the DDR would be required for the recruitment of CDK12 to DNA-damaged sites. Chemical inhibitors of either ATM or ATR did not prevent mCherry-CDK12 localization to laser-irradiated DNA (Fig. 4a,b). In addition, recruitment of CDK12 was preserved in H2AX-KO cells (Fig. 4c and Extended Data Fig. 20a,b). On the other hand, inhibition of PARP, which has also been implicated in the control of DDR and transcription at damaged sites^55^, prevented CDK12 recruitment to DDR-sites (Fig. 4d,e). PARP1 recruitment to DDR sites was independent of either ATR or ATM activity, suggesting that PARP1 and ATM/ATR may control DDR signaling independently (Extended Data Fig. 20c). We next evaluated whether ATM, PARP or CDK12 inhibition, would affect RNA synthesis at the damaged MS2-reporter in I-SceI cells. Inhibition of either PARP or CDK12 rescued transcription, while their combined inhibition led to a marginal increase of the percentage of cells showing transcriptional rescue, consistent with the evidence that PARP and CDK12 are on the same signaling axis (Fig. 4f). As previously reported^56^, ATM inhibition rescued transcription (Fig. 4f). Combination of ATM and CDK12 inhibition did not significantly increase the rescue of transcription, compared to the single inhibitors (Fig. 4f). This suggests that while CDK12 recruitment to damaged sites is ATM-independent, the ATM and PARP/CDK12 signaling may converge downstream to control transcription at damaged sites.

**Figure 4.**
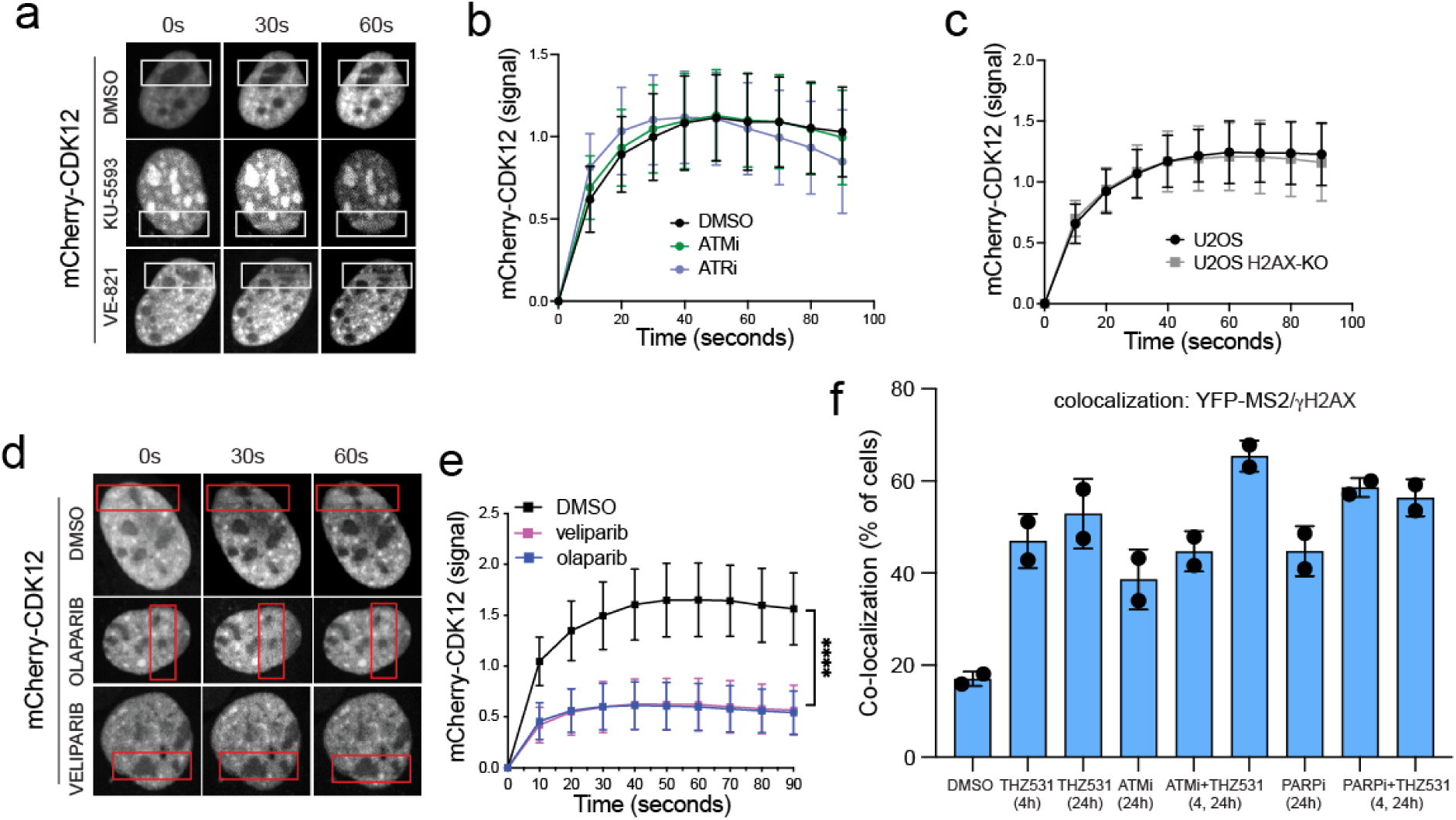
PARP dependent DDR-signaling is necessary for CDK12 recruitment to DNA-damaged sites. **a,b**, Recruitment of mCherry-CDK12 to laser-damaged sites in U2OS cells treated with ATMi (10 µM KU-5593) or ATRi (10 µM VE-821). (**a**) Irradiated areas are highlighted by a white frame. DMSO: n=14; ATMi: n=17; ATRi: n=13. **c**, Recruitment of mCherry-CDK12 to laser-damaged sites in H2AX knock-out cells. U2OS wt: n=18; U2OS KO: n=26. **d**,**e**, Recruitment of mCherry-CDK12 to laser-damaged sites in U2OS cells treated with 1 µM olaparib or 10 µM veliparib. (**d**) Irradiated areas are highlighted by a red frame. DMSO: n=6; veliparib: n=27; olaparib: n=25. **f**, Nascent RNA analysis at the DNA-damaged TRE-MS2 reporter locus of U2OS-TRE-I-SceI-19 cells treated with the indicated compounds. DMSO: n=124, n=130; THZ531: 4h n=144, n=130; THZ531: 24h n=150, n=146; ATMi: 24 n=132, n=115; ATMi 4h + THZ531: 24h n=129, n=101; PARPi: n=146, n=167; PARPi 4h + THZ531: 24h n=126, n=154.

### CDK12 prevents the recruitment of CDK9 to damaged genes

Our experiments showed that CDK12 recruitment to damaged promoter-proximal regions depended upon the recruitment of active transcriptional complexes (tTA-ER) and the presence of a transcribing RNA polymerase. This possibly implies that on these loci CDK12 may repress the activity of the elongating RNAPII. Given the prominent role of CDK9 in elongation, we tested whether CDK12 might modulate CDK9 recruitment to damaged DNA. Coherently with transcriptional inhibition, CDK9 did not localize at laser-irradiated DNA in mock-silenced cells. Instead, silencing of CDK12 allowed the recruitment of CDK9 on damaged DNA (Fig. 5a, Extended Data Fig.21a and Supplementary Video 4,5). Similarly, I-SceI-induced DSBs reduced the co- localization of CDK9 and mCherry-tTA-ER; conversely, silencing of CDK12 engendered CDK9 positive spots proximal to DDR foci (DH2AX) and transcriptional foci (mCherry-tTA-ER) (Fig. 5b,c and Extended Data Fig.21b). This suggested that CDK12 blocks the transcription of damaged genes by preventing CDK9 recruitment.

**Figure 5.**
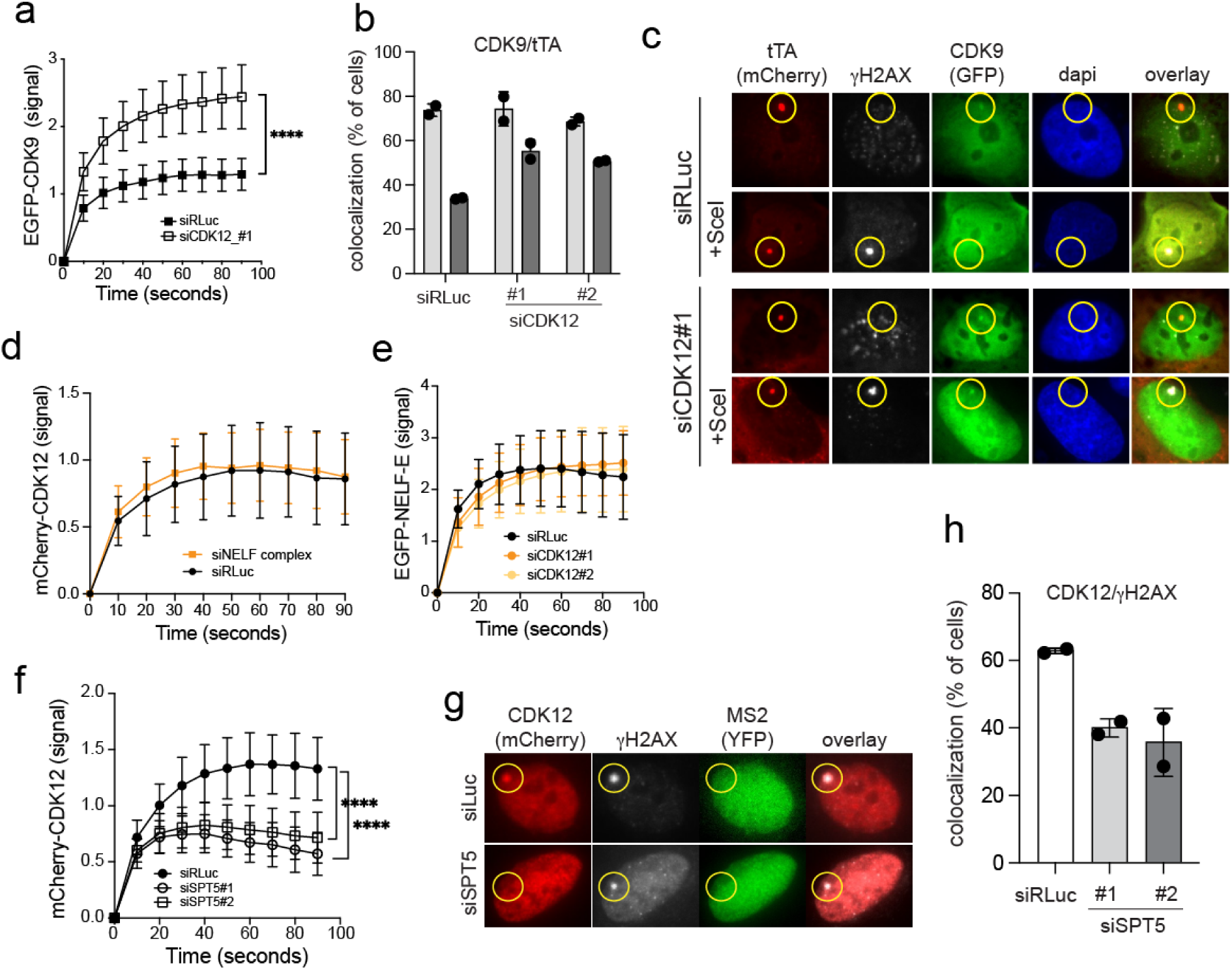
CDK12 is recruited by SPT5 to prevent CDK9 loading to DNA-damaged sites. **a**, Silencing of CDK12 allows the recruitment of EGFP-CDK9 to laser-damaged DNA. siRLuc: n=23; siCDK12#1: n=21. **b**,**c,** Colocalization of mCherry-tTA-ER and EGFP-CDK9 at the promoter of the MS2 reporter in U2OS-TRE-I-SceI-19 cells, at either un-damaged promoters (mock) or I-SceI cut promoters (+SceI). Loss of CDK12 expression enhances localization of CDK9 at the DNA damaged reporter. (**b**) quantification of two independent experiments. siRLuc: n=70, n=150; siCDK12#1: n=83, n=122; siCDK12#2: n=67, n=122; siRLuc+SceI: n=136, n=166; siCDK12#1+SceI: n=90, n=107; siCDK12#2+SceI: n=103, n=148. **d**, Recruitment of mCherry-CDK12 at laser-damaged DNA in NELFc silenced. Silencing the NELF complex (NELFc) does not affect CDK12 recruitment to laser-damaged DNA. siRLuc: n=13; siNELF complex: n=18. **e**, NELF-E recruitment at laser-damaged DNA in CDK12 silenced cells. siRLuc: n=7; siCDK12#1: n=10; siCDK12#2: n=18. **f**, Recruitment of mCherry-CDK12 at laser-damaged DNA in SPT5 silenced cells. siSPT5 prevents the recruitment of CDK12 to laser-damaged loci. siRLuc: n=18; siSPT5#1: n=21; siSPT5#2: n=19. **g**,**h,** Loss of mCherry-CDK12 localization at the I-SceI cut reporter locus of U2OS-TRE-I-SceI-19 cells, following SPT5 silencing. (**g**), representative images. (**h**), bar plot of the fraction of cells showing colocalization of mCherry-CDK12 and DH2AX. siRLuc: n=123, n=135; siSPT5#1: n=126, n=136; siSPT5#2: n=118, n=119.

Two major complexes, NELFc and DSIF, regulate pause-release and elongation by controlling CDK9 recruitment to RNAPII. The NELF-E subunit of NELFc can repress transcription at damaged promoters^57^, thus raising the question of whether NELFc may mediate CDK12 association to damaged loci. However, silencing of NELF-E or other NELFc subunits did not affect CDK12 recruitment on micro-irradiated nuclear regions (Fig. 5d and Extended Data Fig. 22a-c). Moreover, silencing of CDK12 did not prevent NELF-E recruitment, thus indicating that both proteins are recruited independently to DSBs (Fig. 5e). We next addressed the DSIF complex, a heterodimer composed of SPT4/5. While silencing SPT4 did not alter CDK12 recruitment to DSBs (Extended Data Fig. 22d,e), knock-down of SPT5 prevented it (Fig. 7f and Supplementary Video 6). This was also confirmed in the MS2-reporter line, where silencing of SPT5 decreased the colocalization of CDK12 with DDR foci (Fig. 5g,h). Overall, these results indicate that upon DNA breaks proximal to promoter regions, CDK12 is recruited by the DSIF complex, thus preventing CDK9 loading and processive elongation by RNAPII.

### Loss of CDK12 exacerbates replicative stress and transcription-replication conflicts in MYC-overexpressing cells

Replicative stress induced by MYC has been associated to premature initiation of DNA synthesis and the collision of the replicative apparatus with loci transcribed during early S-phase^19,21^. For this reason, we evaluated whether silencing of CDK12 would exacerbate MYC-induced RS. Cell cycle analysis of asynchronous cultures showed that silencing of CDK12 in MYC-overexpressing cells led to an increase in the S-phase fraction (Extended Data Fig. 4c), which, associated with the increased DDR (Fig. 1e), was suggestive of RS. This was confirmed in experiments where cells were released from a mitotic block to allow their synchronous initiation of DNA replication. While MYC stimulated S-phase entry, silencing of CDK12 alone had little effect on cell cycle progression (Fig. 6a). On the other hand, the combination of MYC activation and CDK12 silencing delayed progression through S-phase, as evidenced by EdU incorporation profiles that were still predominantly composed of early S-phase cells (i.e., EdU positive cells with DNA content closer to 2N), both at 12 and 18 hours post-release (Fig. 6a and Extended Data Fig. 23a). Overall, this suggested that loss of CDK12 increased MYC-induced RS. This was supported by quantitative analysis of DNA replication foci by PLA: combined MYC activation and CDK12 loss increased the number of PCNA/EdU foci, indicating an increased compensatory firing of replication origins, which is typically observed upon RS (Extended Data Fig. 23 b,c). Consistent with the onset of RS, the increased DH2AX observed upon loss of CDK12 and MYC activation was associated with newly replicated DNA (Fig. 6b,c). Given the predominant role of transcription-replication conflicts (TRCs) as a potential source of DSBs positioned near transcribed genes, and considering that our data implicates CDK12 in the control of transcription at damaged genes, we asked whether loss of CDK12 would promote the formation of RNA-DNA hybrids. In particular, we reasoned that the loss of CDK12 might increase the chances of forming RNA-DNA hybrids (and R-loops), due to the lack of transcription inhibition at TRC loci and increased annealing of the nascent RNA on to the template DNA. Indeed, in cells depleted of CDK12, we did observe a robust formation of RNA-DNA hybrids following laser-induced DNA-damage, assessed by enhanced recruitment of RNaseH1- GFP (Fig. 6d,e). For a quantitative assessment of TRCs induced by MYC activation, we employed PLA for the detection of RNA-DNA hybrids (detected by the S9.6 antibody) and newly synthesized DNA (EdU-labelled). Knockdown of CDK12 marginally increased the number of S9.6/EdU foci, while MYC activation increased the formation of S9.6/EdU foci (Fig. 6f,g), as expected given its ability to induce TRCs^19^. Importantly, combined activation of MYC and silencing of CDK12 led to a stronger increase of S9.6/EdU foci, thus indicating that loss of CDK12 may indeed exacerbate TRCs (Fig. 6f,g). While these data suggest increased TRCs, whether or not these RNA-DNA hybrids are causal and or linked to increased R-loop formation will need further assessment. Enhanced Myc-induced TRCs were also confirmed by PLA analysis of PCNA and RNAPII, which showed increased signals upon CDK12 silencing (Extended Data Fig. 23e,f). In line with previous reports^33^, activation of Myc led to decreased PCNA/RNAPII signals, whose significance will need further investigation. The evidence that the silencing of CDK12 triggered MYC-induced TRCs suggested a protective role of CDK12, which might be recruited to TRCs loci, once DNA is damaged. In line with this hypothesis, PLA experiments showed that MycER activation enhanced colocalization of CDK12 with DDR foci (Fig. 6h and Extended Data Fig. 23d).

**Figure 6.**
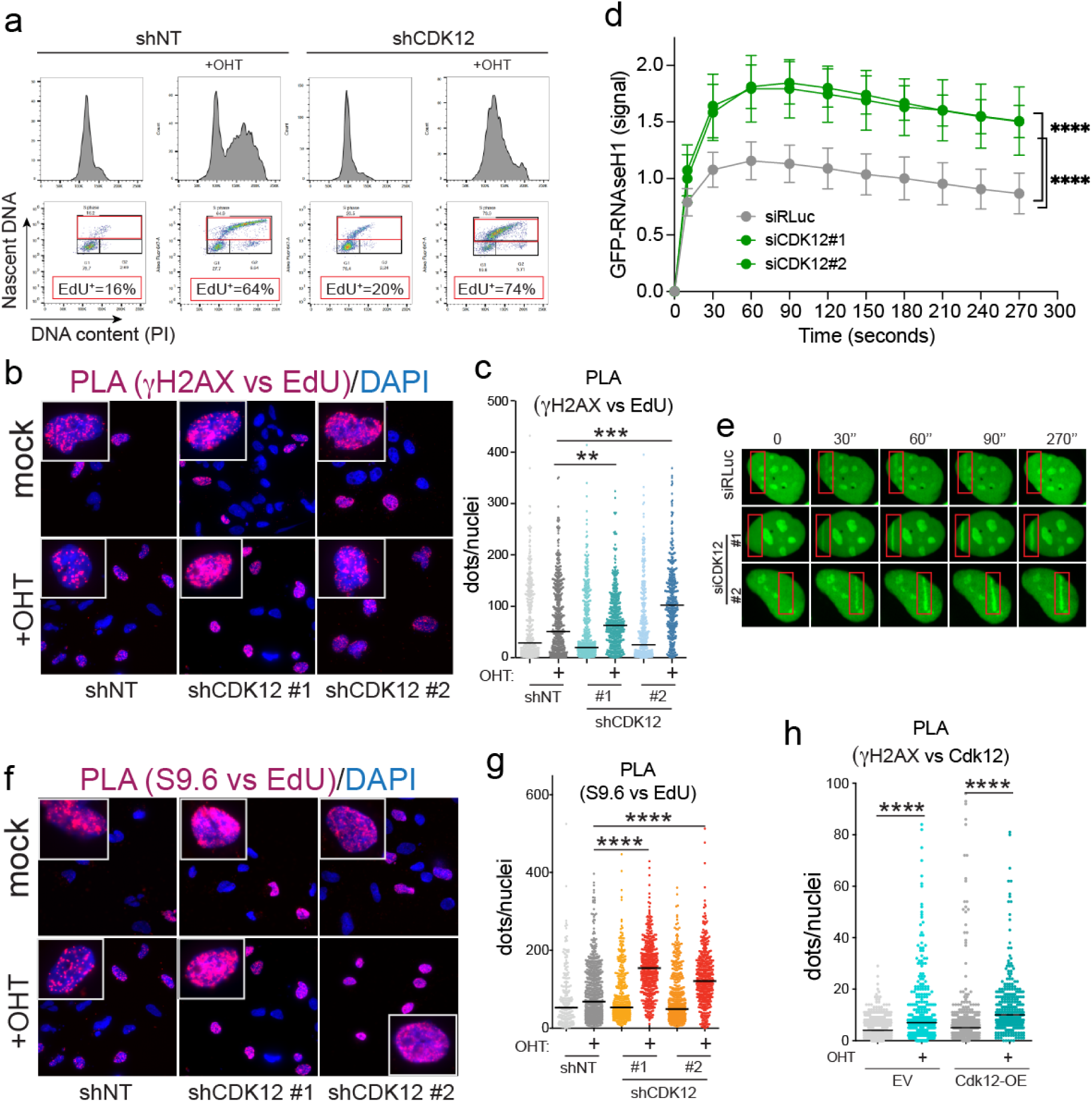
Loss of CDK12 enhances MYC-induced replicative stress. **a**, Cell cycle entry (12 hours post-mitotic release) of U2OS-MycER cells analyzed by FACS. **b**,**c** PLA to evaluate proximity of DH2AX and nascent DNA (EdU-labeled) upon MycER activation (+OHT) and CDK12 silencing. (**b**) representative images. (**c**) bee swarm plot. shNT: n=437 (mock), n=597 (+OHT), shCdk12#1: n=617 (mock), n=567 (+OHT), shCDK12#2: n=438 (mock), n=438 (+OHT). **d**,**e**, Live microscopy analysis of RNAseH1 recruitment to micro-irradiated DNA in U2OS cells silenced for CDK12. **d**, time-series graph. **e**, representative images. siRLuc n=10; siCDK12#1 n=12; siCDK12#2 n=12. **f**,**g,** PLA to evaluate the proximity of RNA-DNA hybrids, stained with the S9.6 antibody and nascent DNA (EdU labelled). (**f**) representative images (**g**) bee swarm plot. shNT: n=211 (mock), 640 (+OHT), shCdk12#1: n=404 (mock), n=445 (+OHT), shCDK12#2: n=538 (mock), n=429 (+OHT). **h,** Bee swarm plot of PLA of CDK12 and DH2AX in U2OS-MycER cells. Where indicated (Cdk12-OE), cells were transfected with a plasmid encoding CDK12. EV: n=534 (mock), n=326 (+OHT), Cdk12-OE: n=410 (mock), n=333 (+OHT)

**Figure 7.**
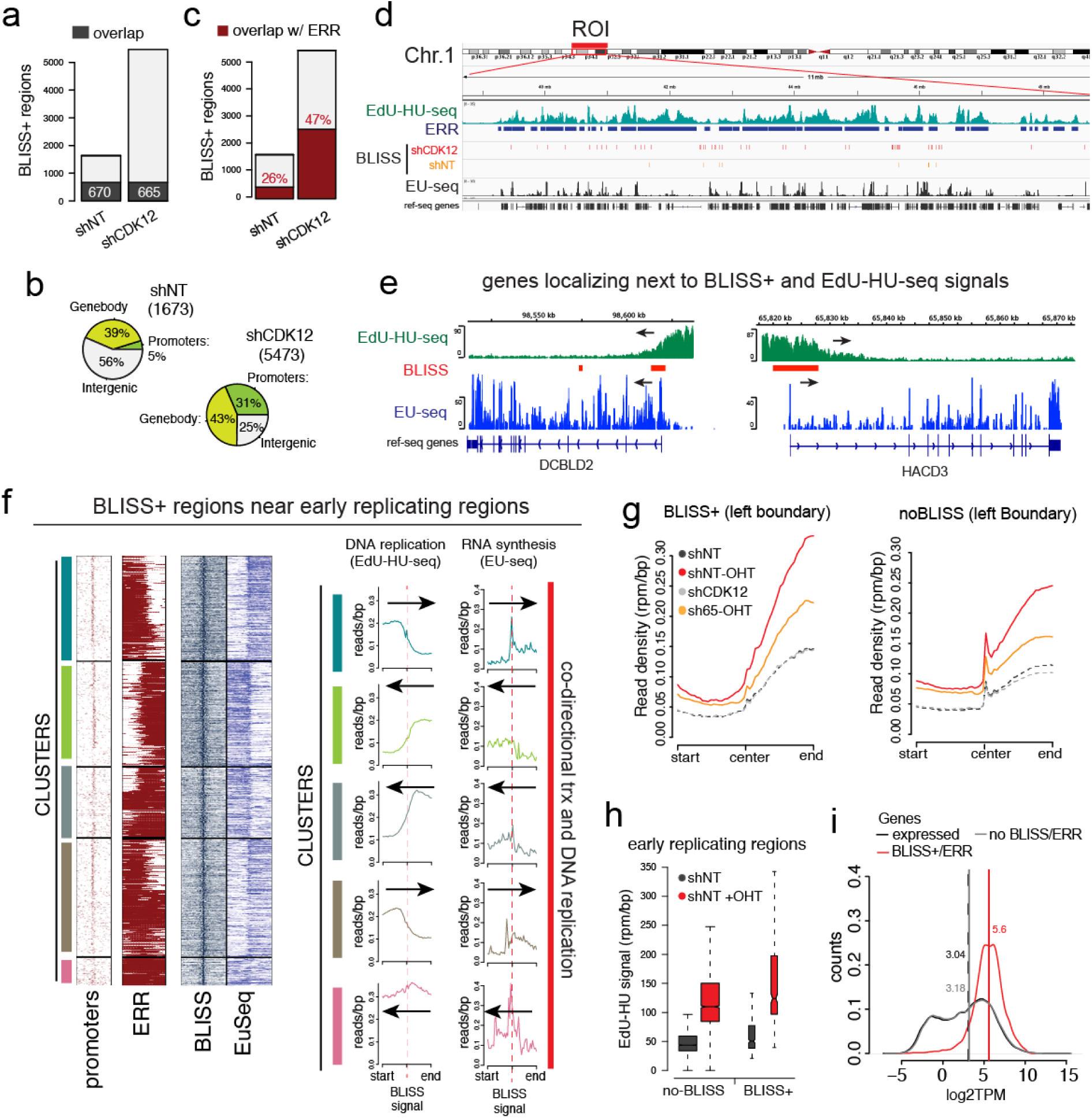
Genome-wide mapping of transcription replication conflicts. **a**, BLISS+ regions identified upon MycER activation and CDK12 (shCDK12) or mock silencing (shNT). In dark gray is the fraction of BLISS+ regions identified in both the shNT and shCDK12 datasets. **b**, Pie charts of the genomic distribution of BLISS+ regions.**c**, Bar plot of the BLISS+ regions identified upon MycER activation and the fraction of BLISS+ regions overlapping with early replicating regions (ERR, indicated in brown). **d**, Genome browser snapshot of the indicated chromosome 1 region (ROI) showing ERR, EdU- HU-seq and EU-seq signals (for MycER activated, shCDK12 cells), and BLISS+ regions (MycER activated cells). **e**, Genome browser snapshots of representative genes identified next to a BLISS+ region and a ERR. Arrows indicate the direction of DNA replication (EdU-HU-seq) and transcription (EU-seq). **f**, Left, clustered heatmaps of BLISS+ regions overlapping with ERR. Heatmaps show promoter position along with EdU-HU-seq (DNA replication), BLISS+ regions (DSBs) and EU-seq signals of 10kb genomic ranges centered on the BLISS signal. The red bar highlights theclusters showing codirectional transcription and DNA synthesis. Right, signal distribution profiles of the regions of each cluster. Arrows indicate the direction of DNA synthesis (assessed by EdU-HU seq) and RNA synthesis (by EU-seq). **g**, EdU-HU-seq signal profiles of the left boundary of ERRs overlapping with a BLISS+ region (left) or ERRs not overlapping with a BLISS+ region (right). **h**, Box plot of the EdU-HU-seq signal shown in (g). **i**, RNA synthesis (EU-signal distribution) of genes next to ERR adjacent to BLISS+ regions (red line), all the expressed genes (black line) or the expressed genes adjacent to an ERR not associated to a BLISS+ region (gray line). Vertical bars indicate the median.

### Loss of CDK12 exacerbates DSBs due to codirectional TRCs at early-replicating regions

The data above suggested TRCs as a cause of DNA damage proximal to newly synthetized DNA. To seek direct evidence, we mapped early replicating regions by EdU-HU-seq^19^ and DSBs by BLISS^58,59^. Since BLISS signals are inherently sparse and “digital”, due to their single-base resolution, we devised an algorithm to identify genomic regions more prone to undergo DSBs (i.e. enriched in BLISS signals), henceforth dubbed as BLISS+ regions.

Of note, while BLISS signals could be detected in any of the conditions considered, BLISS+ regions could only be identified in cells when MycER was activated, possibly reflecting the stochastic nature of DSBs arising in either wild-type or shCDK12 cells. Activation of MycER in shCDK12 cells triggered a four-fold increase in the number of BLISS+ regions (5473 vs 1637, Fig. 7a). Of these, only a minority (12%) was detected also in shNT-MycER cells (Fig. 7a). In addition, BLISS+ regions detected in shCDK12 cells were more conserved among replicates than those detected in wild-type cells (Extended Data Fig. 24a), suggesting that these were recurrent hotspots for DSBs. Also, while BLISS+ regions detected upon MycER activation in shNT-cells were predominantly distributed in either intergenic regions, in shCDK12 cells, BLISS+ regions were preferentially mapped at either promoters or intragenic regions (74%, Fig. 7b). This suggested that transcription was a causal factor in the generation of DSBs when MYC was activated and CDK12 silenced. Given the role of early DNA replication as a potential source for DNA-damage in MYC-overexpressing cells^19^, we used EdU-HU-seq to identify early replicating regions (ERRs, genomic regions enriched in early replicating origins). In shCDK12-MycER cells, half of the BLISS+ regions overlapped with ERR (47%) (Fig. 7c and Extended Data Fig. 24b), either at boundaries or within- ERRs (Fig. 7d and Extended Data Fig. 25a), while the remaining were proximal to sparse EdU signals (Extended Data Fig. 26). ERR-associated BLISS+ regions had two peculiar topological features: (i) they were often positioned at the boundary of the EdU-HU-seq signal, (either on the left or on the right side) and (ii) they were adjacent to the promoter of an expressed gene (Fig. 7e; Extended Data Fig. 25b and 26). Directionality analysis of both DNA and RNA synthesis, indicated that these events were predominantly co-directional, so that almost invariably, BLISS+ regions were positioned between the front of an incoming early DNA replicating locus and the promoter of an expressed gene (Fig. 7e,f and Extended Data Fig. 26). In line with the above, the majority of the BLISS+ regions outside ERR (about 56%) were also associated with sparse EdU-HU-seq signals (due to DNA synthesis from interspersed early replicating origins) and transcribed genes, mostly in a co-directional head to tail orientation of the upcoming DNA replication front and the transcribed gene (Extended Data Fig. 27). On the other hand, in shNT-MycER cells, only 26% of the BLISS+ regions were near ERR (Fig. 7c), and, when proximal to early replicated DNA, they were less frequently associated with transcribed genes (56% of the BLISS+ regions did not overlap with expressed genes) (Fig. 7b). In addition, these BLISS+ regions were neither strongly association with promoters, nor with the other topological features found in shCDK12-MycER cells, since only a minority were positioned between codirectional ERR and transcribed genes (Extended Data Fig. 28, 29). The observation that the DSB-prone regions in shCDK12-MycER cells were associated with expressed genes strongly suggested that loss of CDK12 exacerbated MYC-driven TRCs, leading to accumulation of unresolved DSBs. We next asked whether, at these loci, DNA synthesis and/or RNA expression were modulated by either MYC activation or silencing of CDK12. EdU-HU- seq indicated that DSBs-prone regions in shCDK12-MycER cells (BLISS+ regions) were the replicating regions the most stimulated by MycER activation (Fig. 7g, Extended Data Fig. 30). We also noticed that silencing of CDK12 led to a general reduction of MYC-induced DNA replication since the EdU incorporation levels were lower than those assessed upon MycER activation in mock silenced cells (Fig. 7g,h, Extended Data Fig. 30). This potentially reflected the activity, in trans, of the replication checkpoint. Silencing of CDK12 (alone) did not affect early-replication (Fig. 7g, Extended Data Fig. 30). In addition, while transcripts near BLISS+ regions were not preferentially regulated by MycER activation or by CDK12 silencing, they were synthesized more rapidly than all the other expressed genes (Fig. 7i). Thus, these DSBs were generally associated with regions of strong MYC-induced DNA synthesis and high rates of RNA synthesis, suggesting that (i) loss of CDK12 exacerbated TRCs in regions of intense RNA and DNA synthesis and (ii) these conflicts were triggered by MYC due to its ability to anticipate and boost DNA synthesis (Extended Data Fig. 31).

## DISCUSSION

Transcriptional silencing upon DNA damage is a process regulated by signaling pathways activated by the DDR^60^. This process is triggered by multiple DNA lesions, including DSBs. By inhibiting the activity of transcriptional complexes, cells avoid unnecessary synthesis of potentially faulty mRNAs (a source of transcriptional dependent mutagenesis)^61^, prevent molecular crowding by transcriptional complexes and repair factors, and create a molecular scaffold (e.g., R-loops) to facilitate DNA damage resolution. Current models highlight the prominent roles of apical regulators of the DDR, like ATM and PARP1, which control chromatin remodeling, deposition of repressive histone marks and accessibility at damaged sites^56,62^, thus leading to repression of gene transcription. Our data shows for the first time that CDK12 is recruited to transcribed loci upon DNA damage to repress gene’s expression (Extended Data Fig. 32). This implies an active mechanism that allows selective localization of CDK12 at damaged sites. Genetic dissection of potential upstream regulators revealed that CDK12 localization at these sites is independent from the DDR signaling controlled by ATM, and from DH2AX but instead relies on PARP activity. This may suggest a branched control of transcription at damaged sites, whereby ATM regulates the epigenetic state and accessibility^56^ while PARP may have the added function of regulating RNAPII activity by recruiting CDK12. Since several RNAPII associated factors are PARylated following DNA damage^55^, it is possible that these may also promote CDK12 localization, possibly contributing to transcriptional inhibition at damaged loci. Selective recruitment of CDK12 at damaged DNA also required an elongation competent RNAPII, as it was impaired by inhibition of initiation, by triptolide, or inhibition of elongation by DRB.

RNAPII escape from promoters is dynamically regulated by the association with the NELF complex and the DSIF complex, which, by binding to “opposite” sides of the RNAPII holoenzyme control its pausing and pause-release. Two subunits of the NELF complex, NELF-A and NELF-E, are recruited to RNAPII upon DNA damage to repress transcription^57^. Our data suggest that this is independent of CDK12 since (i) silencing of CDK12 or its inhibition did not prevent NELF-E localization to damaged DNA and (ii) silencing of NELF-E or the full complex did not affect CDK12 localization to damaged DNA. Instead, CDK12 recruitment required SPT5, one of the two subunits of the DSIF complex. This suggests an independent control of DNA damage induced pausing by the NELF complex and by the CDK12/DSIF complex.

DSIF, once phosphorylated by pTEFb, is converted into a positive elongation factor stably associated to RNAPII. Therefore, our observation that CDK9 inhibition by DRB suppresses localization CDK12 at damaged and transcribed loci suggests that the “elongating” form of SPT5 favors CDK12 recruitment. This also implies that, at least in principle, CDK12 could also support transcriptional inhibition at intragenic DNA damaged sites, and not only at promoter proximal sites.

CDK12 is recurrently mutated in several cancers: in ovarian and prostate cancers^63,64^ homozygous (loss of function) mutations are associated with genomic instability characterized by tandem focal duplication at gene-dense regions located at early and late replicated domains^63,65,66^. While CDK12 was reported to regulate processing and expression of genes responsible for DDR signaling and homologous recombination, these genes are not altered in CDK12 mutant tumors^63^. In addition, CDK12 loss of function tumors display genomic instability, mutational profiles and therapeutic responses that are different from homology-repair deficient (HRD) tumors^63,64,66,67^, thus indicating that CDK12 mutant tumors are distinct from HRD-tumors and that loss of CDK12 affects genome integrity also by mechanisms distinct from homology directed repair.

In light of our findings, we propose that CDK12, besides regulating processing and expression of HR-genes, may protect genome integrity by repressing gene transcription and thus facilitating the resolution of DNA damage due to TRCs^68^.

CDK12 is also recurrently amplified in tumors of different origin, suggesting its potential duality in cancer as either an oncogene or a tumor suppressor, depending on the context^39,69^. This is similar to other DDR genes, which, depending on the genetic background, may support or suppress tumor growth. Coherently with a protective function of CDK12 in tumors cells, we and others, have found that CDK12 is required to prevent cytotoxic DNA damage and favor cell survival upon oncogenic activation of MYC^47,51^. In particular, we report here that loss of CDK12 predisposes cells to TRCs and RS. Replicative stress has emerged as a hallmark of oncogene driven proliferation which may act as a tumor suppressive barrier or a therapeutic liability^6^. Several lines of evidence indicated that MYC-induced RS is restrained by safeguarding mechanisms that, once activated, lead to accumulation of cytotoxic DNA damage^24,70^. Still to be fully established are the potential causes of RS and the engaged safeguarding pathways^70^. Recent evidence suggests that anticipated S-phase entry driven by MYC may predispose TRCs, thus leading to fork stalling and accumulation of DNA breaks^19^. In line with this, RS triggered by CDK12 loss led to the accumulation of DNA breaks preferentially located between early replicated regions and genes transcribed in a co-directional orientation. Such regions were characterized by strong MYC-driven DNA replication, but were not selective for MYC-regulated genes. This suggests that MYC-induced DNA replication, rather than transcription, is precipitating these TRCs. Based on the evidence presented here, it is likely that loss of CDK12 (which leads to DDR resistant gene expression) may affect how efficiently DNA damage is resolved at these sites. It thus follows that the increased detection of DSBs might be more likely due to defective repair than to increased TRC frequencies. Other possibilities exist: for instance, we cannot exclude that DDR activation at stalled forks (in the absence of DSBs) might activate CDK12 to stop transcription and avoid fork collapse and DSBs.

Several factors have been implicated as causal in RS-induced genomic instability, here the inactivation of CDK12 allowed to map the precise location on DSBs. This suggests that the management of DNA damage at these sites of potential TRCs is a major liability in MYC-driven cancers. On a broader scale, this may imply a prominent role for TRCs as source of genome instability that could be exploited therapeutically to induce cytotoxic DNA damage.

We posit that future advancements in the mechanistic understanding of processes controlling transcription at damaged genes have potential for the identification of valuable therapeutic targets in cancer treatment.

## Limitations of our study

While loss of function analyses have allowed the identification of factors that regulate CDK12 recruitment to damaged genes and the dissection of how CDK12 modulate RNAPII activity, lack of biochemical data and structure-and-function analyses prevents a precise definition of how CDK12 is recruited to damaged genes or a detailed description of how CDK12 may control CDK9 recruitment. Further studies will be needed to fully define this pathway.

## Acknowledgments

This work was in part supported by the AIRC investigator grant IG-13135 to S.C.. R.B. was funded by the Karolinska Institutet KID Funding Program supporting doctoral students. All BLISS experiments were funded through grants by the Swedish Research Council (grant no. 2018-02950) and the Ragnar Söderberg Foundation (Fellows in Medicine 2016) to N.C.

We thank Ramiro Vazquez, Bruna Caridi, Gaia Sambruni, Emanuele Martini, Simona Rodighiero e Chiara Soriani (IEO Imaging Unit), Simona Ronzoni (IEO-FACS facility), Luca Rotta, Thelma Capra e Salvatore Bianchi (IEO-IIT Genomic Unit) for technical assistance; Bruno Amati, Yilli Docksani, Fabrizio D’adda di Fagagna e Giovanni Tonon for critical discussions and suggestions. We thank Y. Akira, S.P. Jackson, R.H. Medema, B. Majello, N. Ayoub, A. Shibata, G.B. Morin and T. Ikura for sharing reagents.

## Author Contributions

L.C. project supervision, experimental design, experimental work and data analyses. S.R. designed and carried out the siRNA screen set-up, genomic experiments and molecular analyses. N.B. designed and carried out experiments, data analyses. A.A., S.S. and E.P. for experimental work. O.C. carried out bioinformatic and genomic analyses. F.R., M.M. performed the high-throughput siRNA screen, A.A. bioinformatic, statistical and image analyses of the siRNA screen data, M.W. supervised the high-throughput siRNA screen. S.B. designed and carried out live-microscopy experiments, D.P. designed and supervised live-microscopy experiments, R.B. (experiments) and N.C. (supervision) for sBLISS experiments. L.C., N.B. and O.C. contributed to manuscript preparation. S.C. study conceptualization, project supervision, data analyses and manuscript preparation. All the Authors revised the manuscript.

## Data availability

Genomic data are deposited in GEO (GEO accession number: GSE236553.

## Code availability

Computing code for reproduction is available upon request.

## References

1 Macheret, M. & Halazonetis, T. D. DNA replication stress as a hallmark of cancer. Annu Rev Pathol 10, 425–448, doi:10.1146/annurev-pathol-012414-040424 (2015).

2 Burrell, R. A. et al. Replication stress links structural and numerical cancer chromosomal instability. Nature 494, 492–496, doi:10.1038/nature11935 (2013).

3 Gaillard, H., Garcia-Muse, T. & Aguilera, A. Replication stress and cancer. Nat Rev Cancer 15, 276–289, doi:10.1038/nrc3916 (2015).

4 Wilhelm, T., Said, M. & Naim, V. DNA Replication Stress and Chromosomal Instability: Dangerous Liaisons. Genes (Basel) 11, doi:10.3390/genes11060642 (2020).

5 Bowry, A., Kelly, R. D. W. & Petermann, E. Hypertranscription and replication stress in cancer. Trends Cancer 7, 863–877, doi:10.1016/j.trecan.2021.04.006 (2021).

6 Halazonetis, T. D., Gorgoulis, V. G. & Bartek, J. An oncogene-induced DNA damage model for cancer development. Science 319, 1352–1355, doi:10.1126/science.1140735 (2008).

7 Baillie, K. E. & Stirling, P. C. Beyond Kinases: Targeting Replication Stress Proteins in Cancer Therapy. Trends Cancer 7, 430–446, doi:10.1016/j.trecan.2020.10.010 (2021).

8 da Costa, A., Chowdhury, D., Shapiro, G. I., D’Andrea, A. D. & Konstantinopoulos, P. A. Targeting replication stress in cancer therapy. Nat Rev Drug Discov 22, 38–58, doi:10.1038/s41573-022-00558-5 (2023).

9 Keller, K. M. et al. Target Actionability Review: a systematic evaluation of replication stress as a therapeutic target for paediatric solid malignancies. Eur J Cancer 162, 107–117, doi:10.1016/j.ejca.2021.11.030 (2022).

10 Schaub, F. X. et al. Pan-cancer Alterations of the MYC Oncogene and Its Proximal Network across the Cancer Genome Atlas. Cell Syst 6, 282–300 e282, doi:10.1016/j.cels.2018.03.003 (2018).

11 Kress, T. R., Sabo, A. & Amati, B. MYC: connecting selective transcriptional control to global RNA production. Nat Rev Cancer 15, 593–607, doi:10.1038/nrc3984 (2015).

12 Sabo, A. et al. Selective transcriptional regulation by Myc in cellular growth control and lymphomagenesis. Nature 511, 488–492, doi:10.1038/nature13537 (2014).

13 Wolf, E., Lin, C. Y., Eilers, M. & Levens, D. L. Taming of the beast: shaping Myc-dependent amplification. Trends Cell Biol 25, 241–248, doi:10.1016/j.tcb.2014.10.006 (2015).

14 Lin, C. Y. et al. Transcriptional amplification in tumor cells with elevated c-Myc. Cell 151, 56–67, doi:10.1016/j.cell.2012.08.026 (2012).

15 Nie, Z. et al. c-Myc is a universal amplifier of expressed genes in lymphocytes and embryonic stem cells. Cell 151, 68–79, doi:10.1016/j.cell.2012.08.033 (2012).

16 Soucek, L. et al. Modelling Myc inhibition as a cancer therapy. Nature 455, 679–683, doi:10.1038/nature07260 (2008).

17 Donati, G. & Amati, B. MYC and therapy resistance in cancer: risks and opportunities. Mol Oncol 16, 3828–3854, doi:10.1002/1878-0261.13319 (2022).

18 Whitfield, J. R., Beaulieu, M. E. & Soucek, L. Strategies to Inhibit Myc and Their Clinical Applicability. Front Cell Dev Biol 5, 10, doi:10.3389/fcell.2017.00010 (2017).

19 Macheret, M. & Halazonetis, T. D. Intragenic origins due to short G1 phases underlie oncogene-induced DNA replication stress. Nature 555, 112–116, doi:10.1038/nature25507 (2018).

20 Dominguez-Sola, D. et al. Non-transcriptional control of DNA replication by c-Myc. Nature 448, 445–451, doi:10.1038/nature05953 (2007).

21 Robinson, K., Asawachaicharn, N., Galloway, D. A. & Grandori, C. c-Myc accelerates S-phase and requires WRN to avoid replication stress. PLoS One 4, e5951, doi:10.1371/journal.pone.0005951 (2009).

22 Murga, M. et al. Exploiting oncogene-induced replicative stress for the selective killing of Myc-driven tumors. Nat Struct Mol Biol 18, 1331–1335, doi:10.1038/nsmb.2189 (2011).

23 Lecona, E. & Fernandez-Capetillo, O. Targeting ATR in cancer. Nat Rev Cancer 18, 586–595, doi:10.1038/s41568-018-0034-3 (2018).

24 Curti, L. & Campaner, S. MYC-Induced Replicative Stress: A Double-Edged Sword for Cancer Development and Treatment. Int J Mol Sci 22, doi:10.3390/ijms22126168 (2021).

25 Wood, M. A., McMahon, S. B. & Cole, M. D. An ATPase/helicase complex is an essential cofactor for oncogenic transformation by c-Myc. Mol Cell 5, 321–330, doi:10.1016/s1097-2765(00)80427-x (2000).

26 Puccetti, M. V., Adams, C. M., Kushinsky, S. & Eischen, C. M. Smarcal1 and Zranb3 Protect Replication Forks from Myc-Induced DNA Replication Stress. Cancer Res 79, 1612–1623, doi:10.1158/0008-5472.CAN-18-2705 (2019).

27 Rohban, S., Cerutti, A., Morelli, M. J., d’Adda di Fagagna, F. & Campaner, S. The cohesin complex prevents Myc-induced replication stress. Cell Death Dis 8, e2956, doi:10.1038/cddis.2017.345 (2017).

28 Grandori, C. et al. Werner syndrome protein limits MYC-induced cellular senescence. Genes Dev 17, 1569–1574, doi:10.1101/gad.1100303 (2003).

29 Sato, M. et al. The UVSSA complex alleviates MYC-driven transcription stress. J Cell Biol 220, doi:10.1083/jcb.201807163 (2021).

30 Roeschert, I. et al. Combined inhibition of Aurora-A and ATR kinases results in regression of MYCN-amplified neuroblastoma. Nature Cancer 2, 312–326, doi:10.1038/s43018-020-00171-8 (2021).

31 Buchel, G. et al. Association with Aurora-A Controls N-MYC-Dependent Promoter Escape and Pause Release of RNA Polymerase II during the Cell Cycle. Cell Rep 21, 3483–3497, doi:10.1016/j.celrep.2017.11.090 (2017).

32 Herold, S. et al. Recruitment of BRCA1 limits MYCN-driven accumulation of stalled RNA polymerase. Nature 567, 545–549, doi:10.1038/s41586-019-1030-9 (2019).

33 Papadopoulos, D. et al. MYCN recruits the nuclear exosome complex to RNA polymerase II to prevent transcription-replication conflicts. Mol Cell 82, 159–176 e112, doi:10.1016/j.molcel.2021.11.002 (2022).

34 Liu, H., Liu, K. & Dong, Z. Targeting CDK12 for Cancer Therapy: Function, Mechanism, and Drug Discovery. Cancer Res 81, 18–26, doi:10.1158/0008-5472.CAN-20-2245 (2021).

35 Fan, Z. et al. CDK13 cooperates with CDK12 to control global RNA polymerase II processivity. Sci Adv 6, doi:10.1126/sciadv.aaz5041 (2020).

36 Blazek, D. et al. The Cyclin K/Cdk12 complex maintains genomic stability via regulation of expression of DNA damage response genes. Genes Dev 25, 2158–2172, doi:10.1101/gad.16962311 (2011).

37 Dubbury, S. J., Boutz, P. L. & Sharp, P. A. CDK12 regulates DNA repair genes by suppressing intronic polyadenylation. Nature 564, 141–145, doi:10.1038/s41586-018-0758-y (2018).

38 Krajewska, M. et al. CDK12 loss in cancer cells affects DNA damage response genes through premature cleavage and polyadenylation. Nat Commun 10, 1757, doi:10.1038/s41467-019-09703-y (2019).

39 Lui, G. Y. L., Grandori, C. & Kemp, C. J. CDK12: an emerging therapeutic target for cancer. J Clin Pathol 71, 957–962, doi:10.1136/jclinpath-2018-205356 (2018).

40 Murphy, D. J. et al. Distinct thresholds govern Myc’s biological output in vivo. Cancer Cell 14, 447–457, doi:10.1016/j.ccr.2008.10.018 (2008).

41 Paulsen, R. D. et al. A genome-wide siRNA screen reveals diverse cellular processes and pathways that mediate genome stability. Mol Cell 35, 228–239, doi:10.1016/j.molcel.2009.06.021 (2009).

42 Zhou, Y., et al. Metascape provides a biologist-oriented resource for the analysis of systems-level datasets. Nat Commun 10, 1523, doi:10.1038/s41467-019-09234-6 (2019).

43 Chipumuro, E. et al. CDK7 inhibition suppresses super-enhancer-linked oncogenic transcription in MYCN-driven cancer. Cell 159, 1126–1139, doi:10.1016/j.cell.2014.10.024 (2014).

44 Koh, C. M. et al. MYC regulates the core pre-mRNA splicing machinery as an essential step in lymphomagenesis. Nature 523, 96–100, doi:10.1038/nature14351 (2015).

45 Hsu, T. Y. et al. The spliceosome is a therapeutic vulnerability in MYC-driven cancer. Nature 525, 384–388, doi:10.1038/nature14985 (2015).

46 Kessler, J. D. et al. A SUMOylation-dependent transcriptional subprogram is required for Myc-driven tumorigenesis. Science 335, 348–353, doi:10.1126/science.1212728 (2012).

47 Toyoshima, M. et al. Functional genomics identifies therapeutic targets for MYC-driven cancer. Proc Natl Acad Sci U S A 109, 9545–9550, doi:10.1073/pnas.1121119109 (2012).

48 Sun, Z., Zhang, Z., Wang, Q. Q. & Liu, J. L. Combined Inactivation of CTPS1 and ATR Is Synthetically Lethal to MYC-Overexpressing Cancer Cells. Cancer Res 82, 1013–1024, doi:10.1158/0008-5472.Can-21-1707 (2022).

49 Cunningham, J. T., Moreno, M. V., Lodi, A., Ronen, S. M. & Ruggero, D. Protein and nucleotide biosynthesis are coupled by a single rate-limiting enzyme, PRPS2, to drive cancer. Cell 157, 1088–1103, doi:10.1016/j.cell.2014.03.052 (2014).

50 Lankes, K. et al. Targeting the ubiquitin-proteasome system in a pancreatic cancer subtype with hyperactive MYC. Mol Oncol 14, 3048–3064, doi:10.1002/1878-0261.12835 (2020).

51 Gwynne, W. D. et al. Cancer-selective metabolic vulnerabilities in MYC-amplified medulloblastoma. Cancer Cell 40, 1488–1502 e1487, doi:10.1016/j.ccell.2022.10.009 (2022).

52 Magnuson, B. et al. CDK12 regulates co-transcriptional splicing and RNA turnover in human cells. iScience 25, 105030, doi:10.1016/j.isci.2022.105030 (2022).

53 Heine, G. F., Horwitz, A. A. & Parvin, J. D. Multiple mechanisms contribute to inhibit transcription in response to DNA damage. J Biol Chem 283, 9555–9561, doi:10.1074/jbc.M707700200 (2008).

54 Janicki, S. M. et al. From silencing to gene expression: real-time analysis in single cells. Cell 116, 683–698, doi:10.1016/s0092-8674(04)00171-0 (2004).

55 Ray Chaudhuri, A. & Nussenzweig, A. The multifaceted roles of PARP1 in DNA repair and chromatin remodelling. Nat Rev Mol Cell Biol 18, 610–621, doi:10.1038/nrm.2017.53 (2017).

56 Shanbhag, N. M., Rafalska-Metcalf, I. U., Balane-Bolivar, C., Janicki, S. M. & Greenberg, R. A. ATM-dependent chromatin changes silence transcription in cis to DNA double-strand breaks. Cell 141, 970–981, doi:10.1016/j.cell.2010.04.038 (2010).

57 Awwad, S. W., Abu-Zhayia, E. R., Guttmann-Raviv, N. & Ayoub, N. NELF-E is recruited to DNA double-strand break sites to promote transcriptional repression and repair. EMBO Rep 18, 745–764, doi:10.15252/embr.201643191 (2017).

58 Yan, W. X. et al. BLISS is a versatile and quantitative method for genome-wide profiling of DNA double-strand breaks. Nat Commun 8, 15058, doi:10.1038/ncomms15058 (2017).

59 Bouwman, B. A. M. et al. Genome-wide detection of DNA double-strand breaks by in-suspension BLISS. Nat Protoc 15, 3894–3941, doi:10.1038/s41596-020-0397-2 (2020).

60 Min, S., Ji, J. H., Heo, Y. & Cho, H. Transcriptional regulation and chromatin dynamics at DNA double-strand breaks. Exp Mol Med 54, 1705–1712, doi:10.1038/s12276-022-00862-5 (2022).

61 Saxowsky, T. T. & Doetsch, P. W. RNA polymerase encounters with DNA damage: transcription-coupled repair or transcriptional mutagenesis? Chem Rev 106, 474–488, doi:10.1021/cr040466q (2006).

62 Shanbhag, N. M. & Greenberg, R. A. Neighborly DISCourse: DNA double strand breaks silence transcription. Cell Cycle 9, 3635–3636, doi:10.4161/cc.9.18.13171 (2010).

63 Wu, Y. M. et al. Inactivation of CDK12 Delineates a Distinct Immunogenic Class of Advanced Prostate Cancer. Cell 173, 1770–1782 e1714, doi:10.1016/j.cell.2018.04.034 (2018).

64 Popova, T. et al. Ovarian Cancers Harboring Inactivating Mutations in CDK12 Display a Distinct Genomic Instability Pattern Characterized by Large Tandem Duplications. Cancer Res 76, 1882–1891, doi:10.1158/0008-5472.CAN-15-2128 (2016).

65 Sørensen, S. G. et al. Pan-cancer association of DNA repair deficiencies with whole-genome mutational patterns. bioRxiv, 2022.2001.2031.478445 (2022).

66 Menghi, F. et al. The Tandem Duplicator Phenotype Is a Prevalent Genome-Wide Cancer Configuration Driven by Distinct Gene Mutations. Cancer Cell 34, 197–210.e195, doi:10.1016/j.ccell.2018.06.008 (2018).

67 Reimers, M. A. et al. Clinical Outcomes in Cyclin-dependent Kinase 12 Mutant Advanced Prostate Cancer. Eur Urol 77, 333–341, doi:10.1016/j.eururo.2019.09.036 (2020).

68 Yang, Y. et al. Large tandem duplications in cancer result from transcription and DNA replication collision. medRxiv, doi:10.1101/2023.05.17.23290140 (2023).

69 Filippone, M. G. et al. CDK12 promotes tumorigenesis but induces vulnerability to therapies inhibiting folate one-carbon metabolism in breast cancer. Nat Commun 13, 2642, doi:10.1038/s41467-022-30375-8 (2022).

70 Rohban, S. & Campaner, S. Myc induced replicative stress response: How to cope with it and exploit it. Biochim Biophys Acta 1849, 517–524, doi:10.1016/j.bbagrm.2014.04.008 (2015).

