## supplementary figures for "CDK12 controls transcription at damaged genes and prevents MYC-induced transcription-replication conflicts"

*Extended data figure 1*

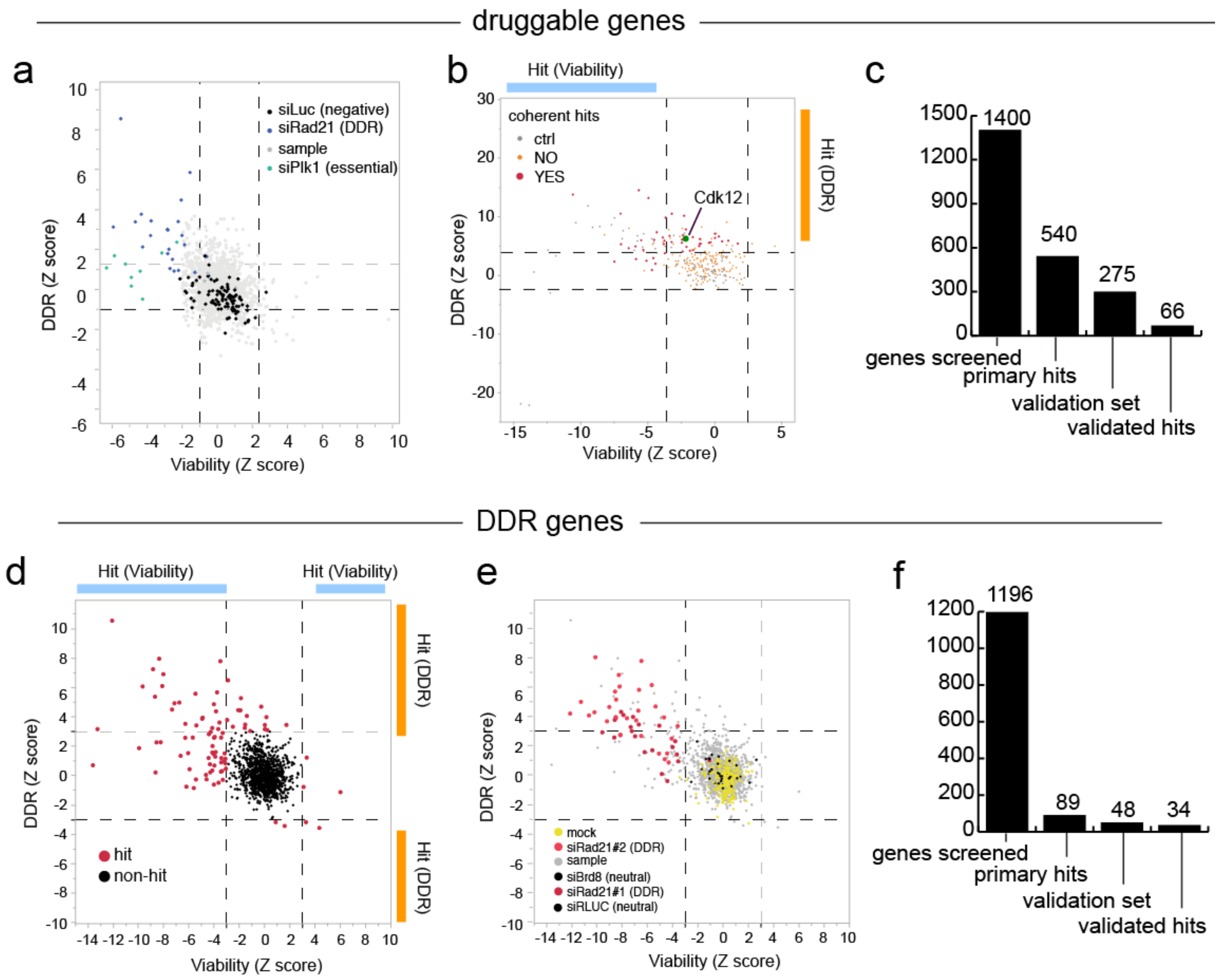


**Extended data figure 1. Summary of the siRNA screens.**

**a**,**b**, Dot plot of the normalized differential DDR score (y axis) and the normalized differential viability (x axis) for each siRNA of the library targeting druggable genes (in gray) and controls. Each dot represents a silenced mRNA. Dashed lines indicate the thresholds used to call the DDR and viability hits. (**a**) primary screen, (**b**) a subset of the secondary screen.

**c**, Bar plot of the genes tested at each stage of the screen of the druggable library.

**d**,**e**, Dot plots of the normalized differential DDR score (y axis) and the normalized differential viability (x axis) for each siRNA of the library targeting druggable genes (primary screen). In (**d**) are highlighted the hits called (shown in red), while in (**e**) are highlighted the controls used to set the thresholds for hit calling.

**f**, Bar plot of the genes tested at each stage of the DDR library screen.

Extended data figure 2

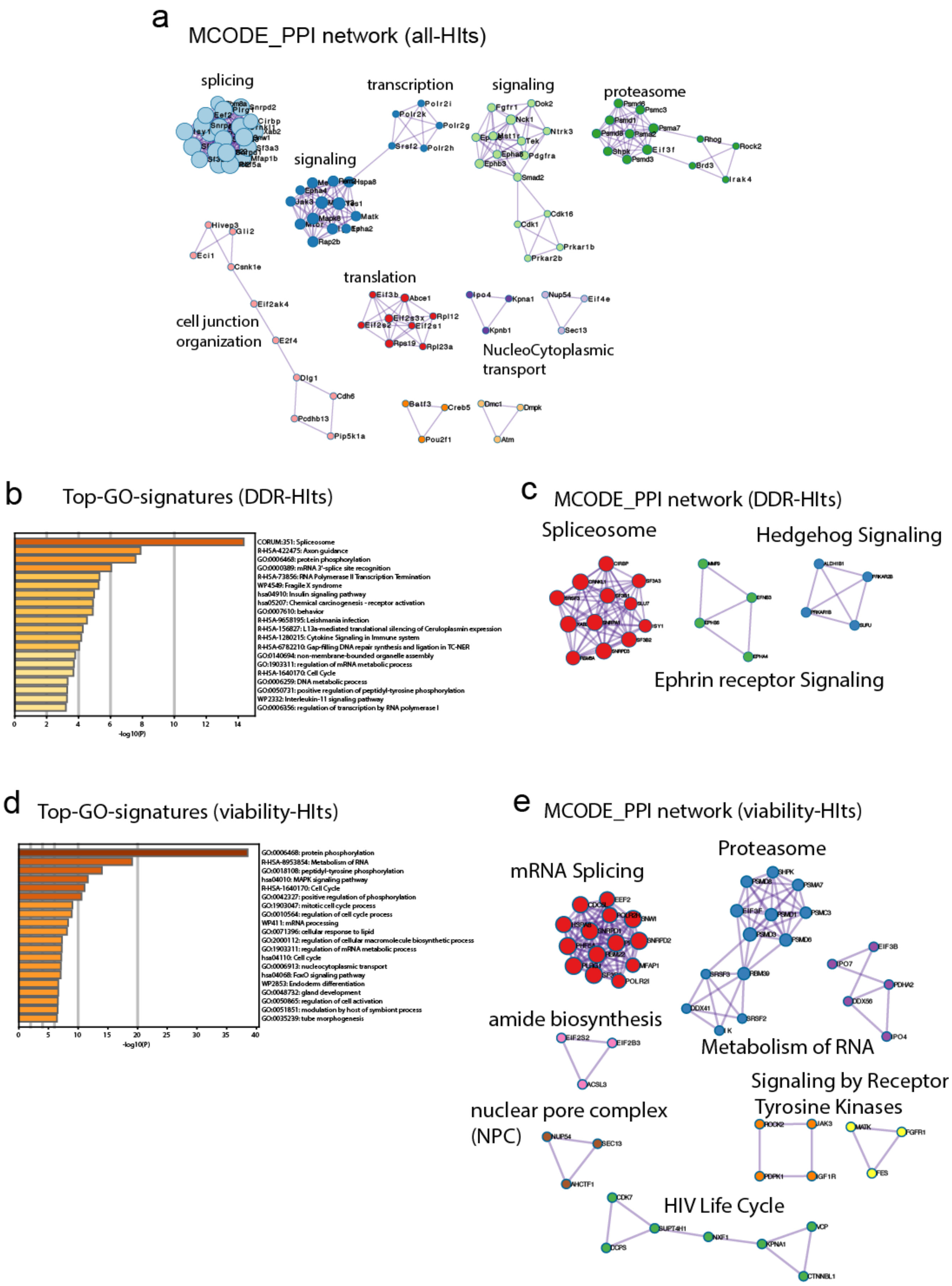


**Extended data figure 2. Metascape analyses of primary hits.**
**a**, MCODE-PPI analysis (enriched protein-protein interaction pathways) of all the hits of the two primary screens. **b**-**e**, Top GO-signatures and MCODE-PPI pathways found in DDR hits (**b**,**c**) and viability hits (**d**,**e**).

*Extended data figure 3*

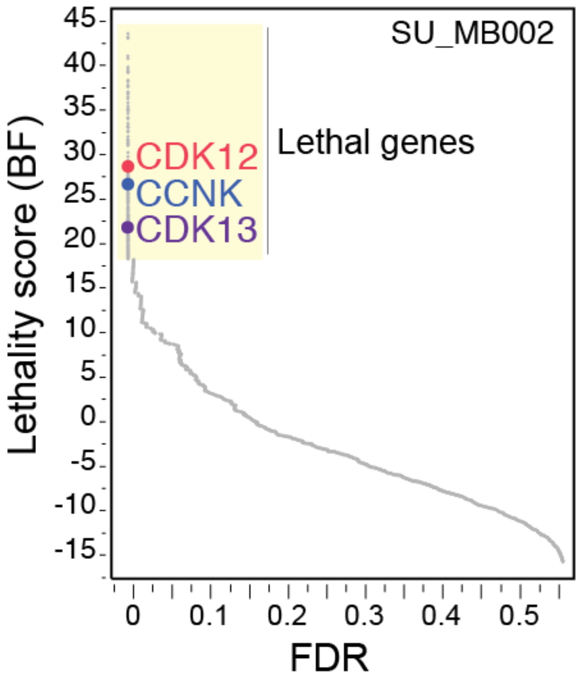


**Extended data figure 3. CDK12, CDK13 and CCNK are synthetic lethal with MYC amplification in Medulloblastoma.**
Waterfall plot of the lethality score of genes silenced in the MYC amplified Medulloblastoma cell line MB002. CDK12, CDK13 and CCNK are among the genes synthetic lethal with MYC amplification. This is a re-analysis of the data reported in Gwynne et al., Cancer Cell 2022.

*Extended data figure 4*

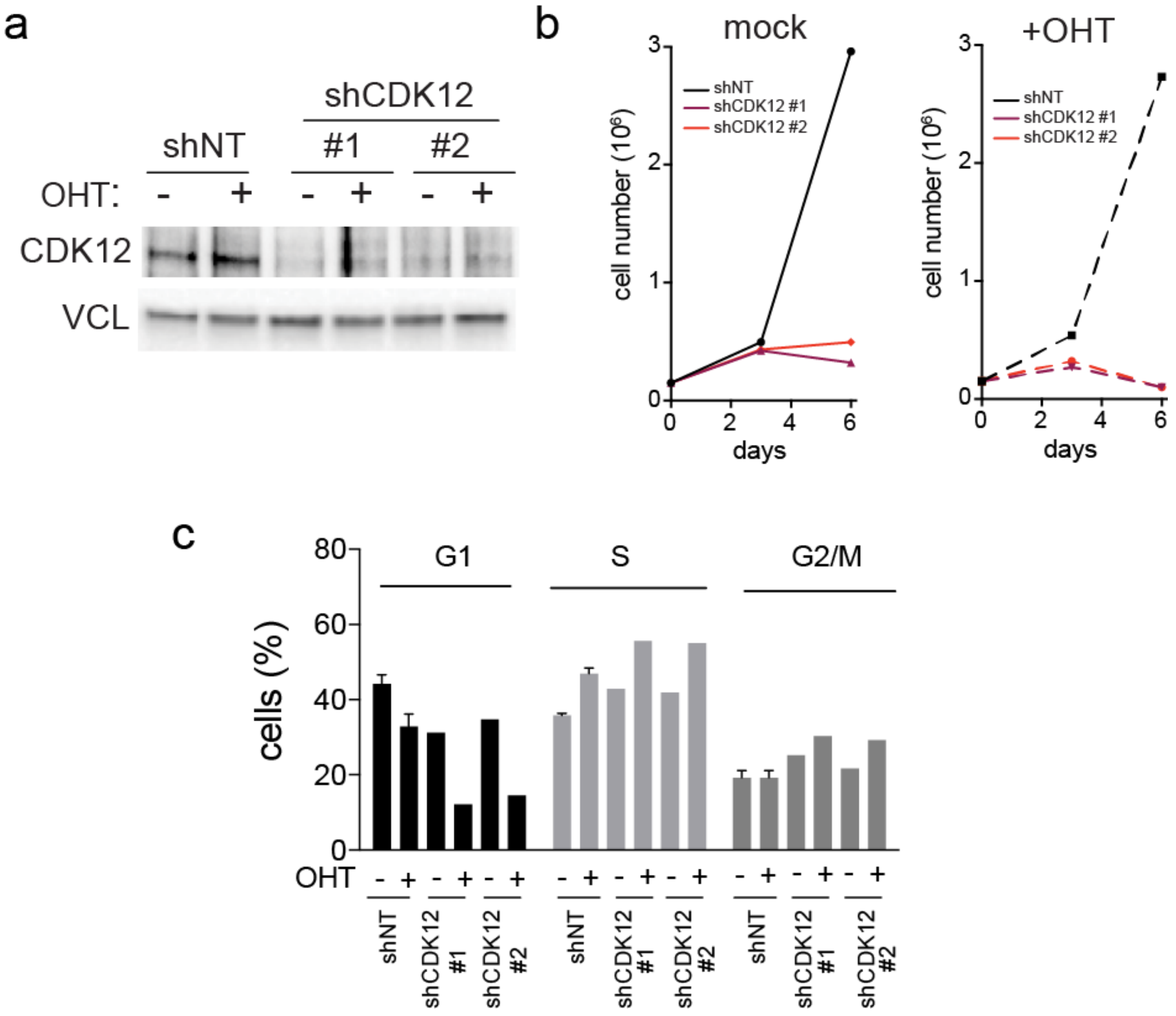


**Extended data figure 4. CDK12 silencing in U2OS-MycER cells.**
**a**, WB analysis of CDK12 levels upon three days of silencing. VCL was used as a loading control.
**b**, Growth curves of U2OS-MycER cells. Left, mock. Right, with activated MycER (+OHT).
**c**, Cell cycle analysis of U2OS-MycER cells, following 48 hours of MycER activation and CDK12 silencing. Bar graphs are the average of triplicates, with standard deviation error bars.

*Extended data figure 5*

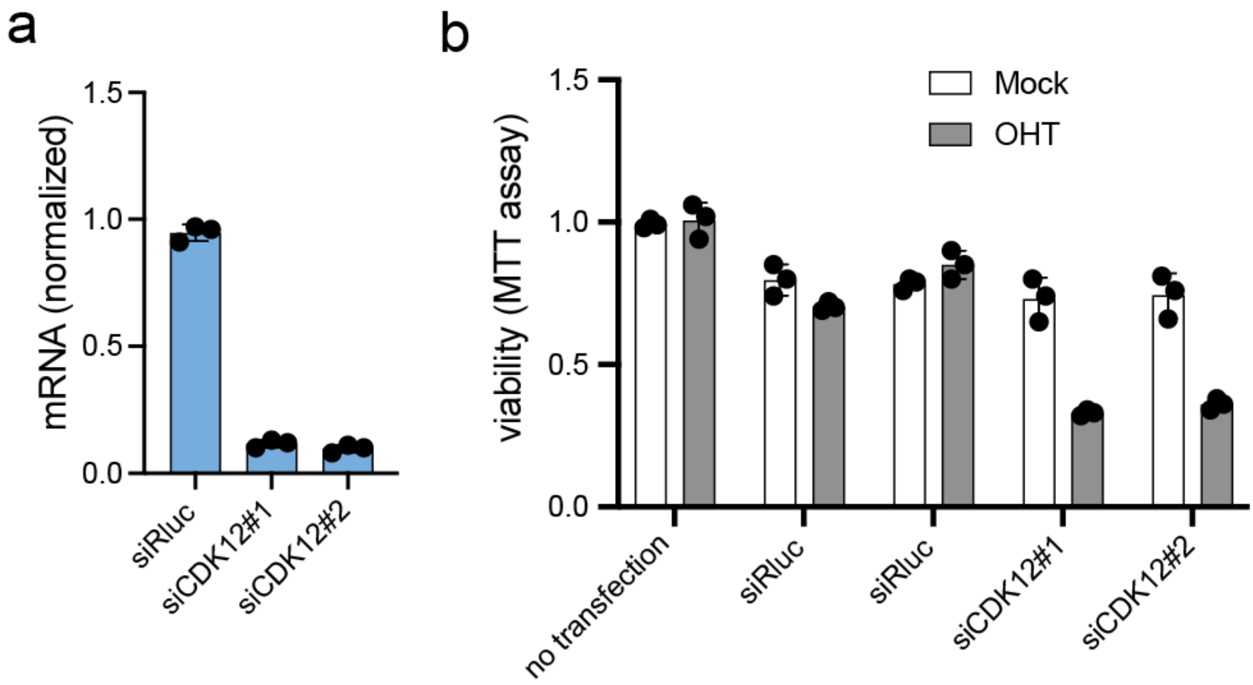


**Extended data figure 5. siRNA mediated silencing of CDK12 impairs viability of MycER-activated cells.**
U2OS MycER cells were analyzed 72 hours post-transfection and activation of MycER (+OHT).
**a**, RT-qPCR analysis. CDK12 values are normalized to RPLP0.
**b**, Normalized cell viability. Bar graphs are the average of triplicates, with standard deviation.

*Extended data figure 6*

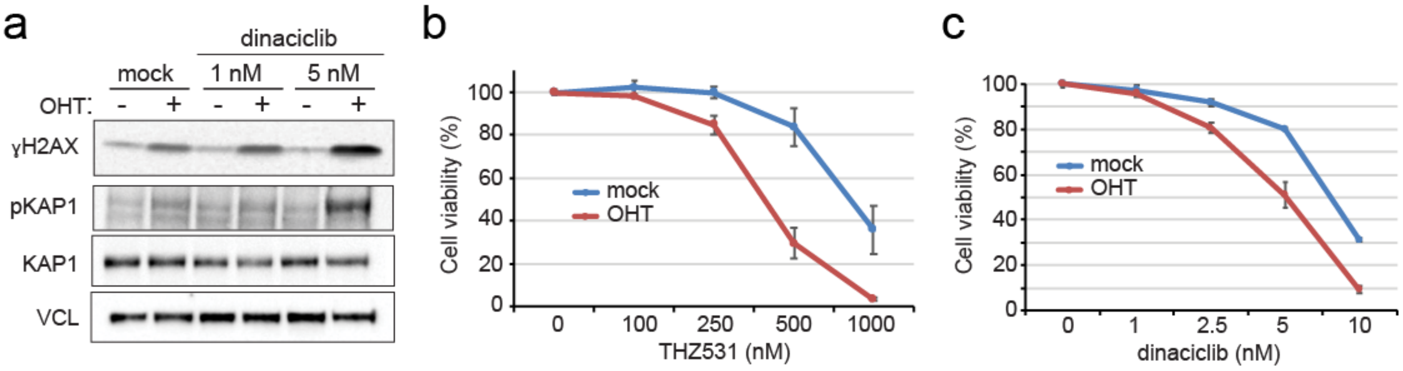


**Extended data figure 6. CDK12 inhibition induces DDR and lethality upon activation of MycER.** U2OS MycER cells were treated with CDK12 inhibitors (THZ531 and dinaciclib). Analyses were performed at 72 hours of drug treatment and activation of MycER (+OHT).

**a**, WB analysis of γH2AX, pKAP1, total KAP1 (DDR marker) and VCL (loading control).

**b**,**c**, Cell viability assessed by CellTiter-Glo assay (Promega). Reported are the average of triplicates with standard deviation.

*Extended data figure 7*

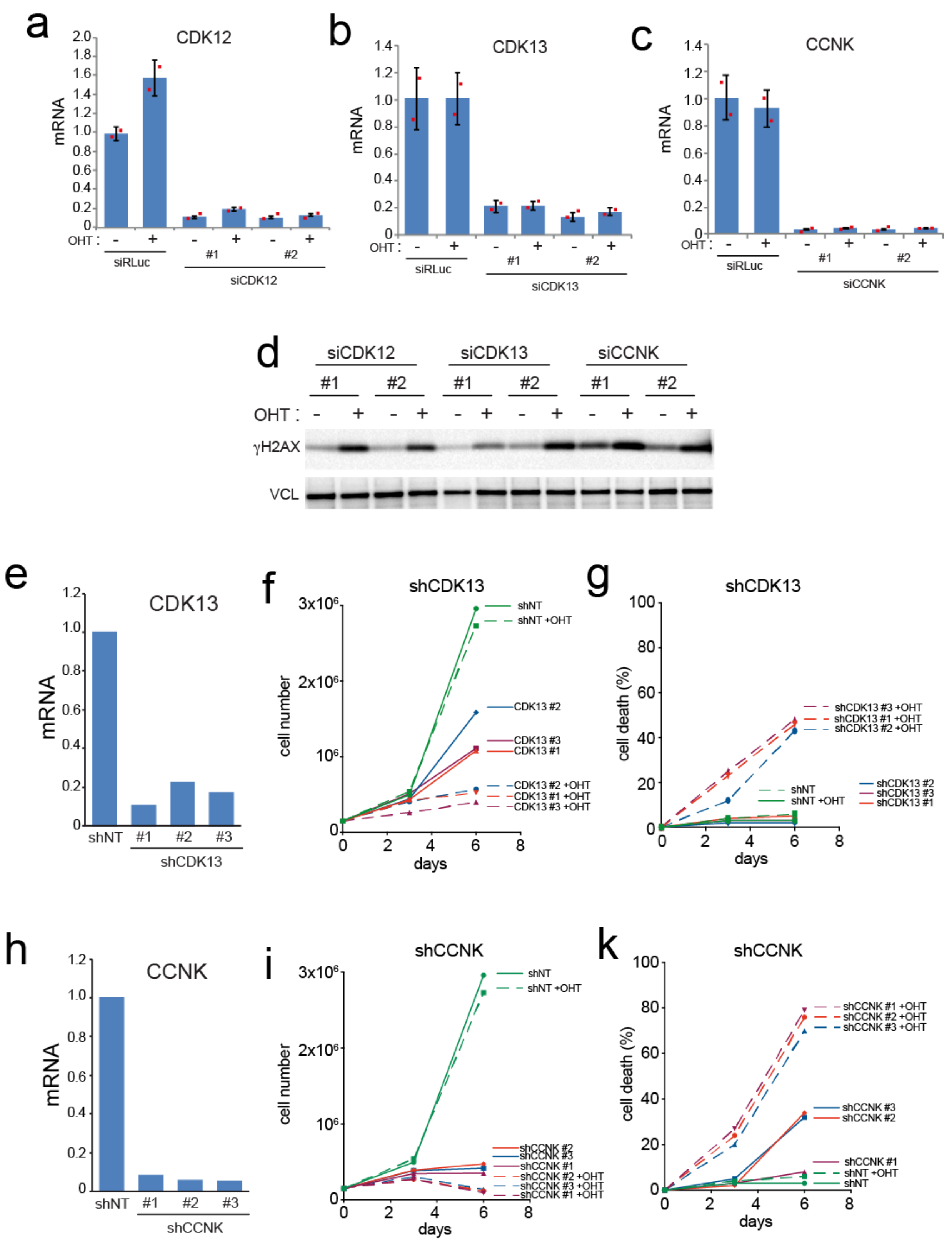


**Extended data figure 7. silencing of CCNK or CDK13 is synthetic lethal with MycER activation.**
**a-c**, RT-qPCR analysis of U2OS-MycER cells transfected with the indicated siRNAs. Values were normalized to RPLP0.
**d**, WB blot analysis of U2OS-MycER cells transfected with the indicated siRNAs. VCL was used as a loading control
**e-g.** Analyses of U2OS-MycER cells infected with lentiviruses encoding shCDK13. (e) RT-qPCR, (f) cell growth curves, (g) cell death by trypan blue assay.
**h-k**, Analyses of U2OS-MycER cells infected with lentiviruses encoding shCCNK. (h) RT-qPCR, (i) cell growth curves, (k) cell death by trypan blue assay.

*Extended data figure 8*

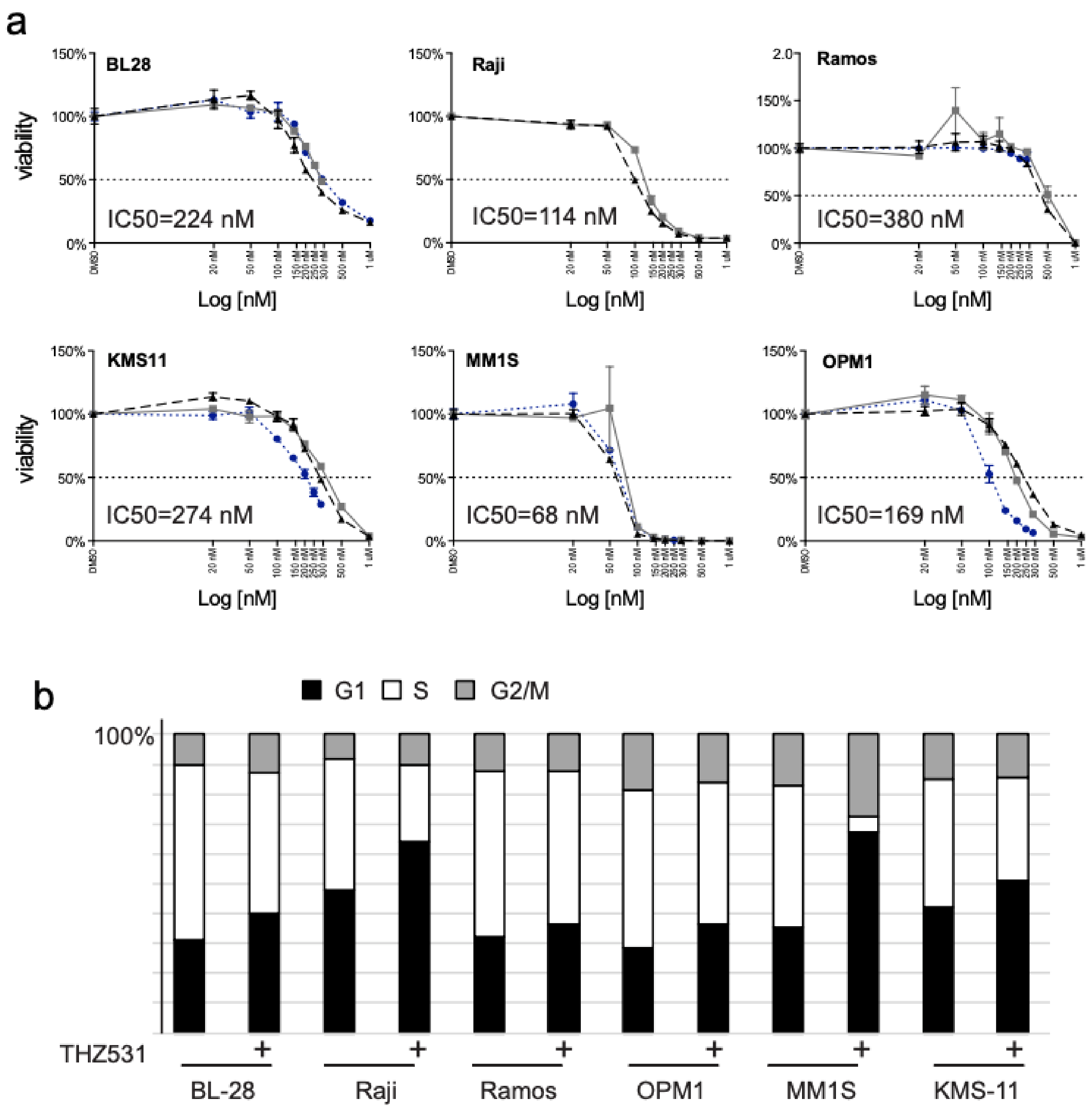


**Extended data figure 8. Pharmacological inhibition of CDK12 in multiple myeloma and Burkitt's lymphoma cell lines.**
**a**, THZ531 dose response curves. Values reported are the average of triplicate measures. Each curve is an independent experiment and IC50s are the average +/- Stdv. Cell viability was assessed by CellTiter-Glo®
**b**, Cell cycle distribution of the indicated cell lines treated with THZ531 at their IC50 concentration. Cell cycle phases were evaluated by BrdU/PI staining and FACS analysis.

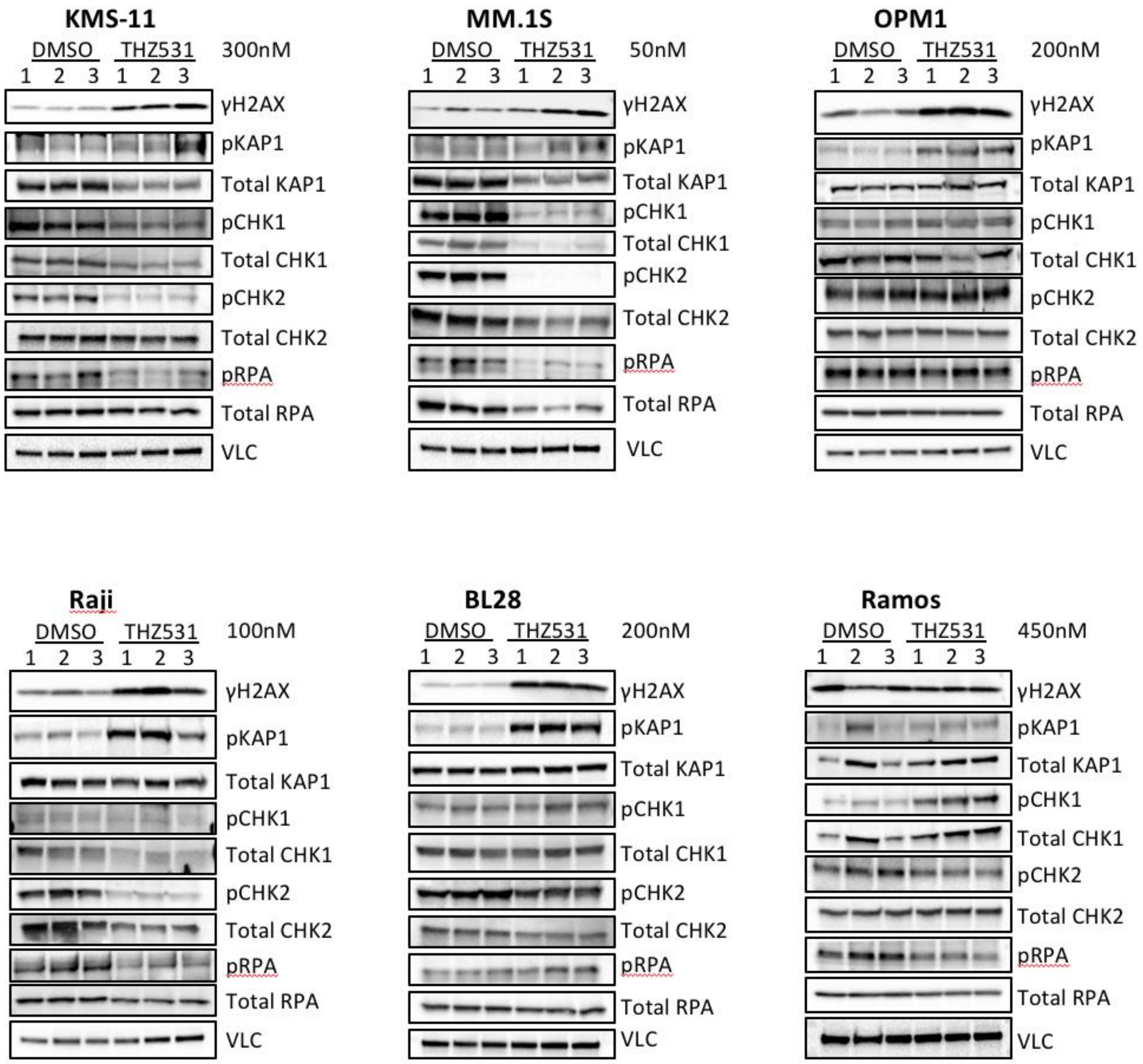


**Extended data figure 9. CDK12 inhibition triggers DDR in MYC-driven multiple myeloma and Burkitt's lymphoma cell lines.**
WB analysis of DDR markers following the treatment of Multiple myeloma (KMS-11, MM.1S and OPM1) and Burkitt's lymphoma (Raji, BL28 and Ramos) cell lines with THZ531 for 48 hours. Vinculin (VCL) was used as loading control.

*Extended data figure 10*

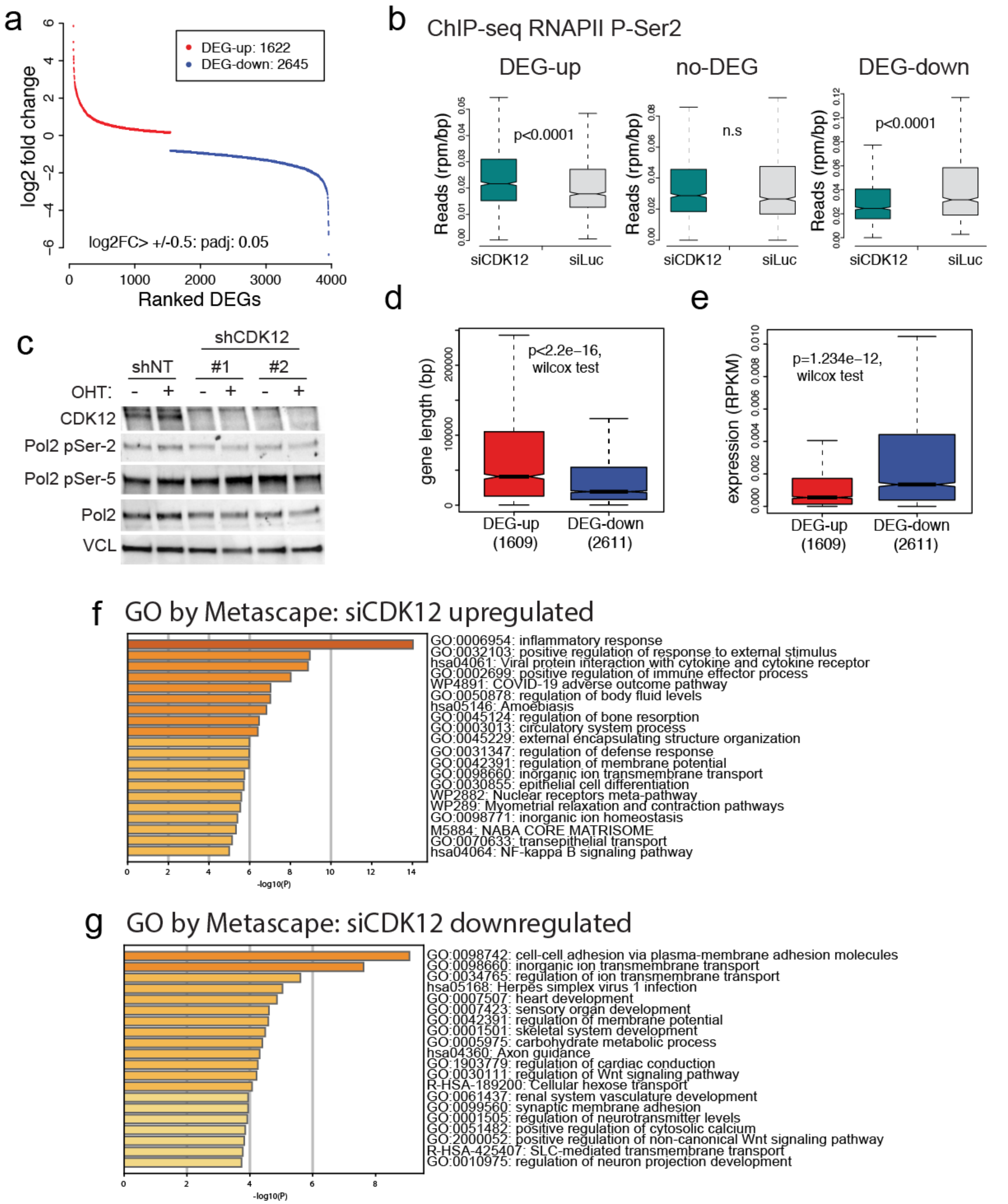


**Extended data figure 10. Genome-wide transcriptional changes upon CDK12 silencing in U2OS cells.** **a**, Differentially expressed genes (DEG) identified by RNAseq analysis in siCDK12 cells. Expression threshold: Log2FC>0.5 or <0.5. Statistical threshold: P.adj (FDR)≤0.5. **b**, Box plot of RNAPII Phospho-Ser-2 ChIP-seq signals on the gene body of DEGs, in either siLuc or siCDK12 U2OS cells. P value=paired t-test
**c**, WB analyses of RNAPII (pol2) levels and phosphorylation upon MycER activation (48hrs) in shNT or shCDK12 U2OS-MycER cells.
**d**,**e**, Box plot of the gene length (**d**) and expression level (**e**) for upregulated (DEG-up) and downregulated (DEG-down) genes.
**f**,**g,** GO analysis of DEGs identified in siCDK12 cells.

*Extended data figure 11*

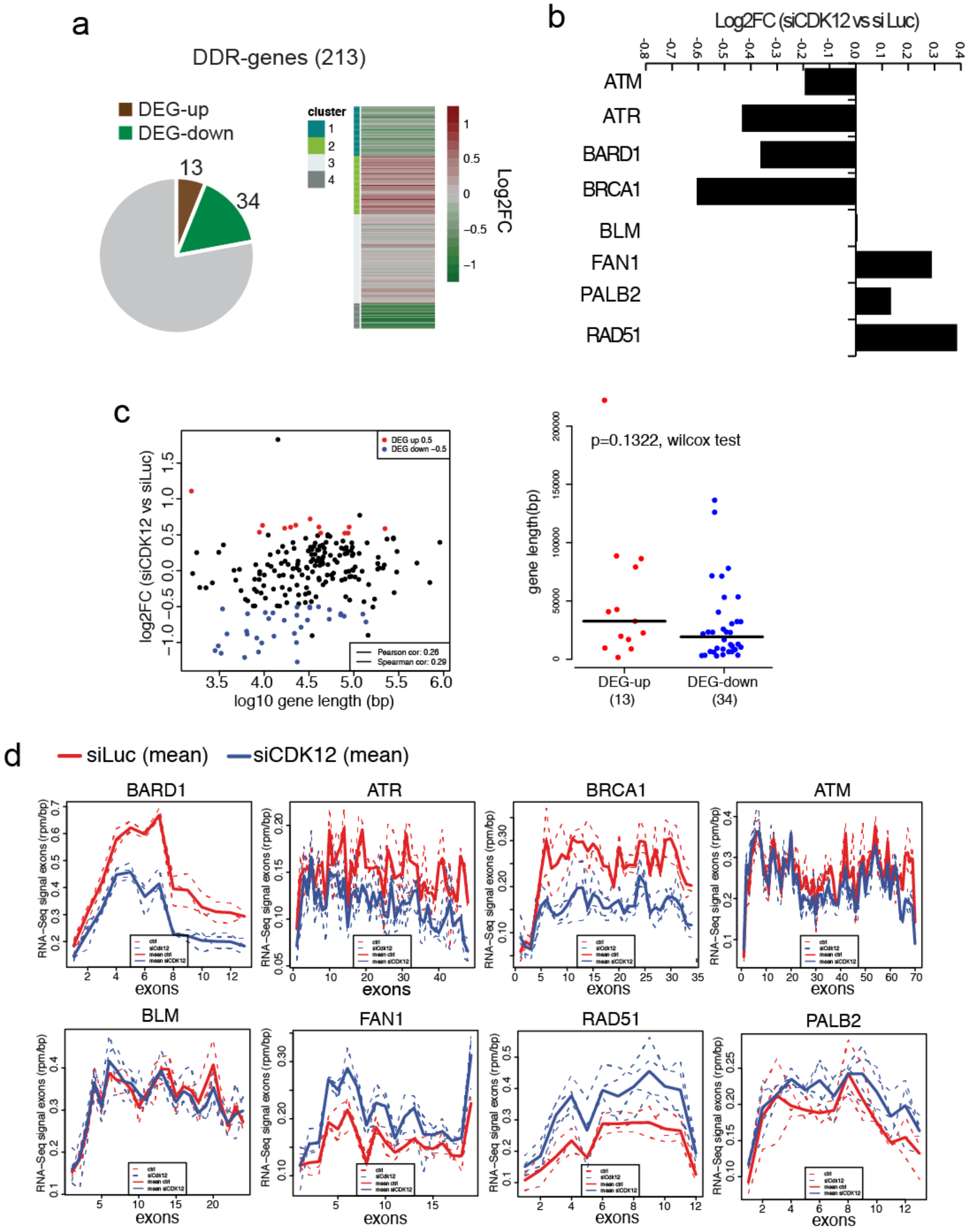


**Extended data figure 11. Expression and processing of DDR genes in siCDK12 U2OS cells. a**, Left, Pie chart of differentially expressed DDR gene. Right, heatmap of relative expression changes (log2FC) of DDR genes following CDK12 silencing. **b**, Differential expression of select DDR genes. **c**, Dot plot (left) and box plot (right) of gene length and expression of DDR genes. This shows a mild positive correlation between gene length and differential expression. **d**, Metaplots of the average exon signals of long DDR genes. Dashed lines are the signal from each RNAseq experiment analyzed. This analysis does not reveal a strong effect of CDK12 silencing on the processing of these DDR genes.

*Extended data figure 12*

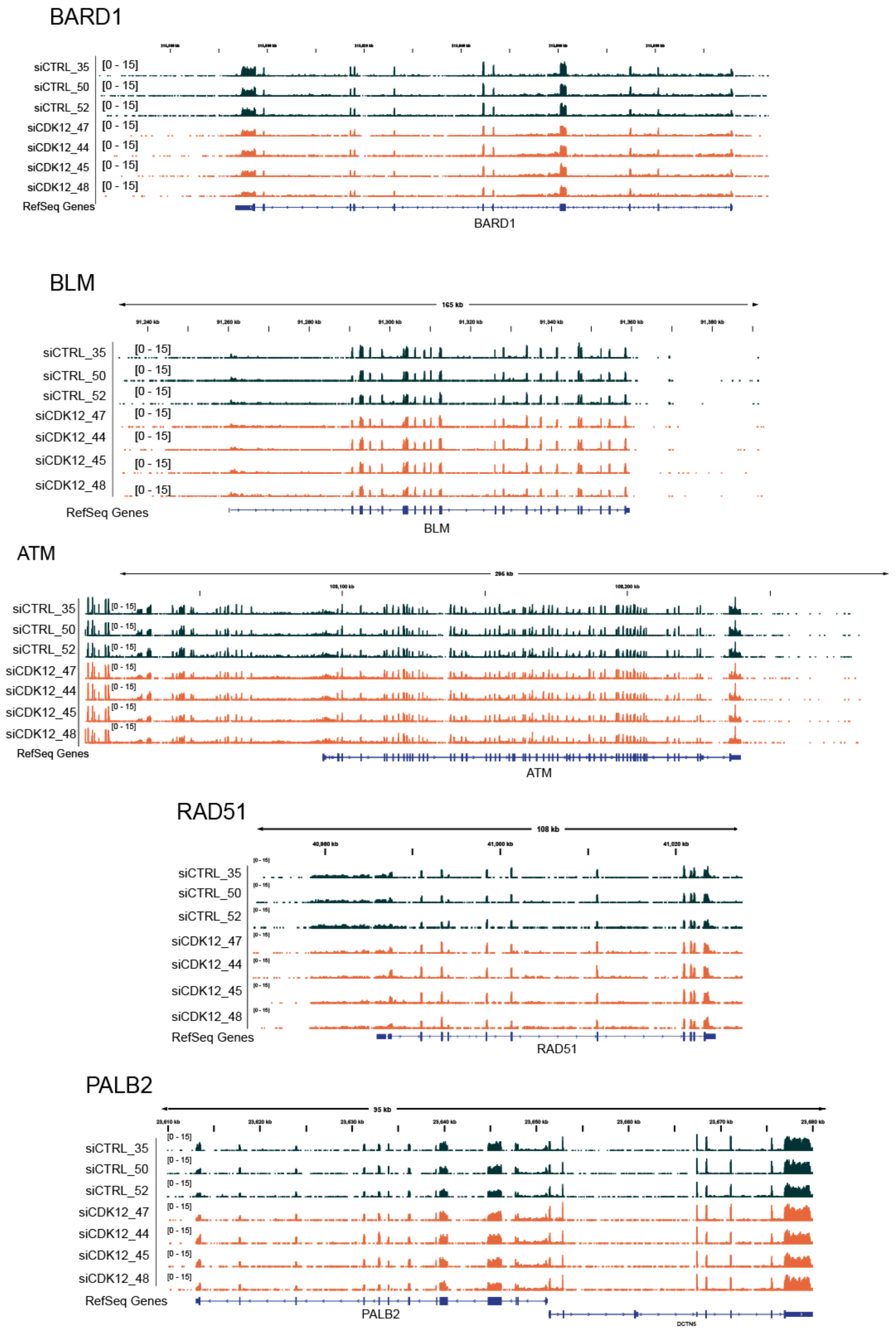


**Extended data figure 12. Lack of early termination or splicing defects in long DDR genes.** Genome browser snapshots of long-DDR genes showing the library normalized RNAseq signals in siCTRL (black) or siCDK12 (orange) U2OS cells.

*Extended data figure 13*

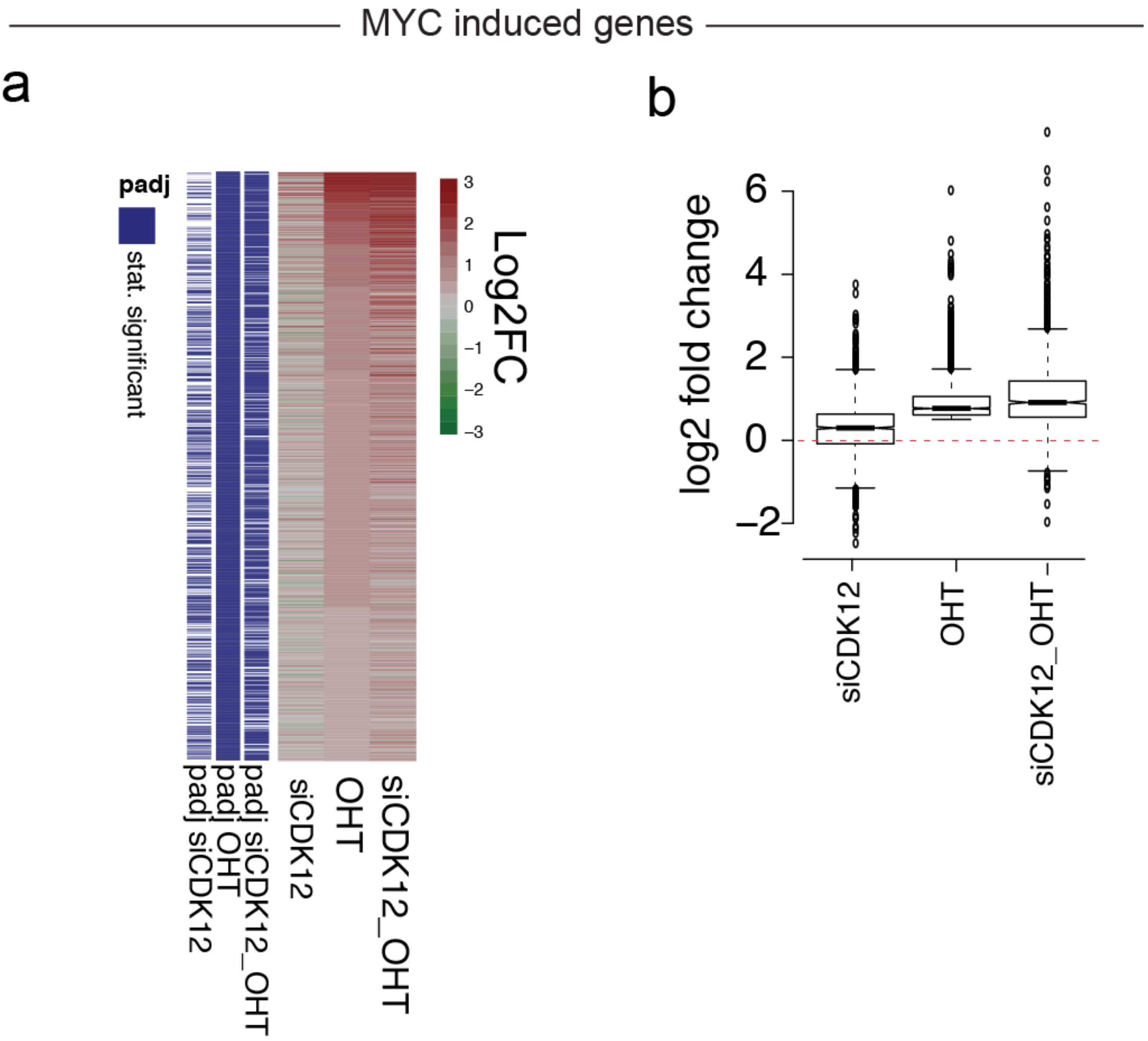


**Extended data figure 13. Analysis of MYC-induced genes upon silencing of CDK12 in U2OS cells.**
**a**, Heatmap of Log2FC of MycER induced genes.
**b**, Box plot of Log2FC of MycER induced genes.

*Extended data figure 14*

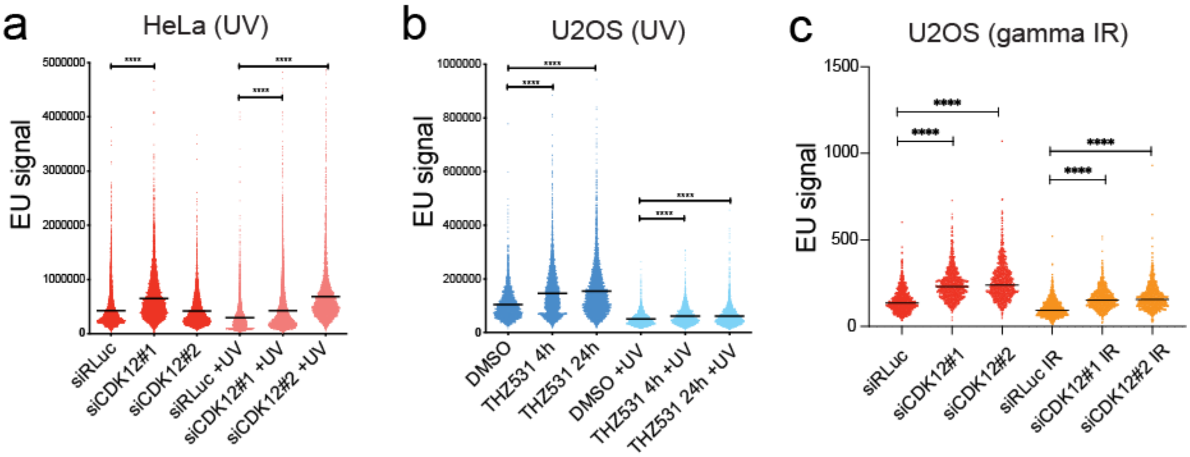


**Extended data figure 14. Nascent RNA synthesis is modulated by CDK12.**
Bee swarm plots of single-cell nascent RNA synthesis (EU incorporation) in mock, UV- or gamma-irradiated cells. Cells were pulsed with 0.5 mM EU 20 minutes before collection. **a**, HeLa cells. siRLuc: n=8200; siCDK12#1: n=8500; siCDK12#2: n=5087; siRLuc+UV: n=5003; siCDK12#1+UV: n=11860; siCDK12#2+UV: n=7065. **b**, U2OS cells. DMSO: n=3180; THZ531 4h: n=5128; THZ531 24h: n=5128; DMSO+UV: n=2367; THZ531 4h +UV: n=2350; THZ531 24h +UV: n=3392.
**c**, U2OS cells. siRLuc: n=1000; siCDK12#1: n=950; siCDK12#2: n=938; siRLuc+UV: n=1057; siCDK12#1+UV: n=888; siCDK12#2+UV: n=987.

*Extended data figure 15*

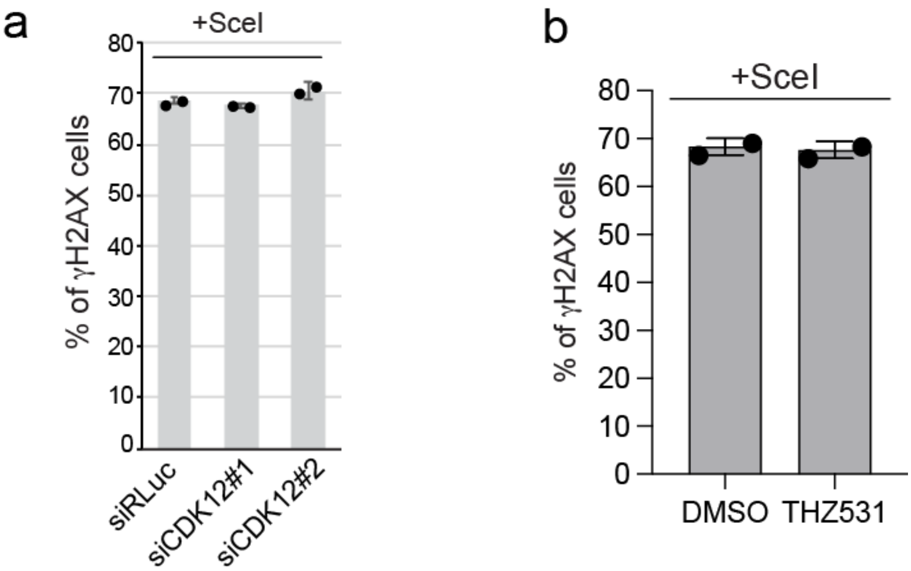


**Extended data figure 15. Efficiency of DSBs induced by I-SceI at the TRE-MS2 reporter in U2OS-TRE-I-SceI-19 cells.**
**a**,**b**, Efficiency of DSBs induced by I-SceI in U2OS-TRE-I-SceI-19 cells upon (**a**) CDK12 silencing or (**b**) its inhibition by THZ531. siRLuc: n=64, n=109; siCDK12#1: n=70, n=102; siCDK12#2: n=67, n=76. DMSO: n=113, n=87; THZ531: n=111, n=96.

*Extended data figure 16*

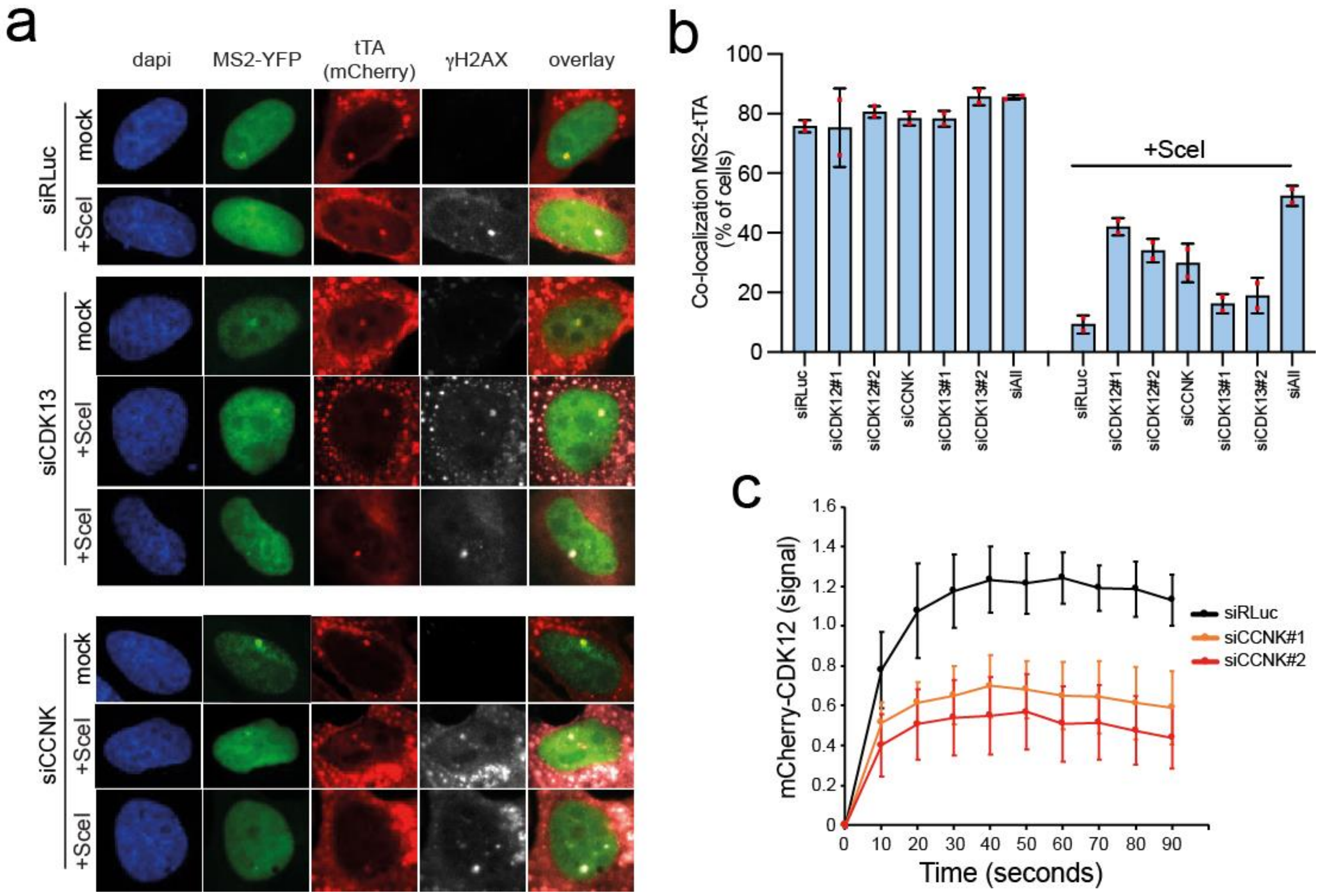


**Extended data figure 16. Silencing of CCNK or CDK13 rescues transcription at the damaged TRE-MS2 reporter.**

**a**,**b**, Representative IF-images (**a**) and bar plot (**b**) of the colocalization of mCherry-tTA-ER, γH2AX and YFP-MS2 signals upon silencing of CDK13 or CCNK in U2OS-TRE-I-SceI-19 cells. Where indicated, cells were transfected with I-SceI (+SceI) to induce DSBs on the TRE-MS2 reporter. Average of two independent experiments. siRLuc: n=105, n=101; siRLuc +SceI: n=113, n=105; siCCNK#1: n=65, n=90; siCCNK#1+SceI: n=75, n=87; siCDK13#1: n=111, n=110; siCDK13#1+SceI: n=110, n=173; siCDK13#2: n=106, n=104; siCDK13#2+SceI: n=136, n=183; siAll (siCDK12, siCDK13, and siCCNK): n=106, n=93; siAll (siCDK12, siCDK13, and siCCNK) +SceI: n=108, n=115;

**c**, Kinetics of recruitment of mCherry-CDK12 to laser-damaged sites in U2OS cells transfected with siCCNK or siLuc (mock). siRLuc: n=9; siCCNK#1: n=4; siCCNK#1: n=9.

Extended data figure 17

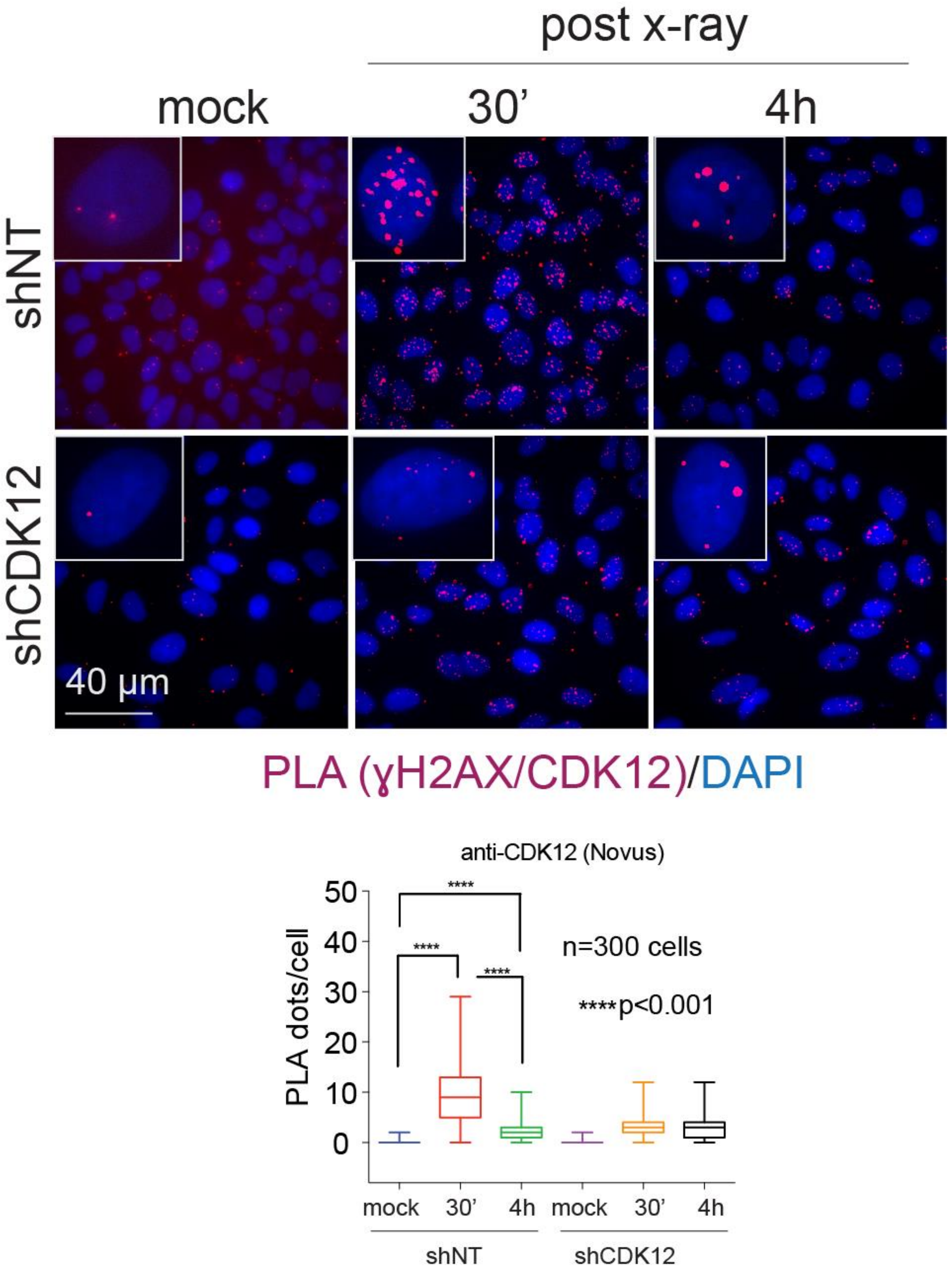


**Extended data figure 17. CDK12 colocalizes with DDR-foci on transcribed loci.**

PLA of γH2AX-CDK12 in X-ray irradiated sh-CDK12-U2OS cells. Top, snapshots. Bottom, box plot. shNT: n= 287(noIR), n= 317(30min), n=275 (4 hours); shCDK12: n= 303(noIR), n= 177(30min), n= 306(4 hours)

*Extended data figure 18*

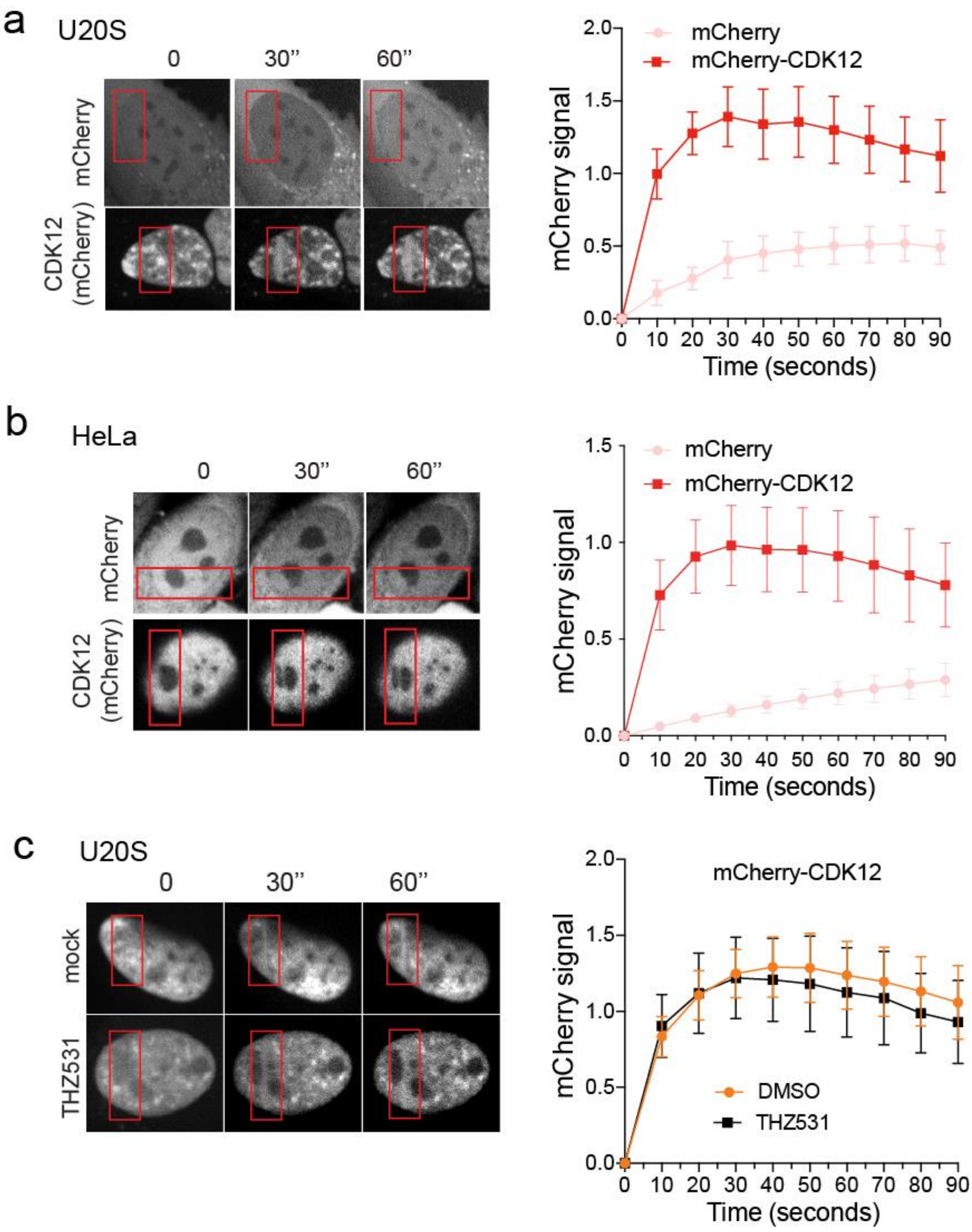


**Extended data figure 18.**

**a**,**b**, Recruitment of mCherry-CDK12 to laser micro-irradiated nuclear areas, assessed in (**a**) U2OS (mCherry: n=12; mCherry-CDK12: n=12) or (**b**) HeLa cells upon CDK12 silencing (mCherry: n=30; mCherry-CDK12: n=12). Left, representative images. Right, time series plot.

**c**, Recruitment of mCherry-CDK12 at laser micro-irradiated nuclear areas in cells treated with the CDK12 inhibitor THZ531 (mCherry: n=7; mCherry-CDK12: n=10). Left, representative images. Right, time series plot.

*Extended data figure 19*

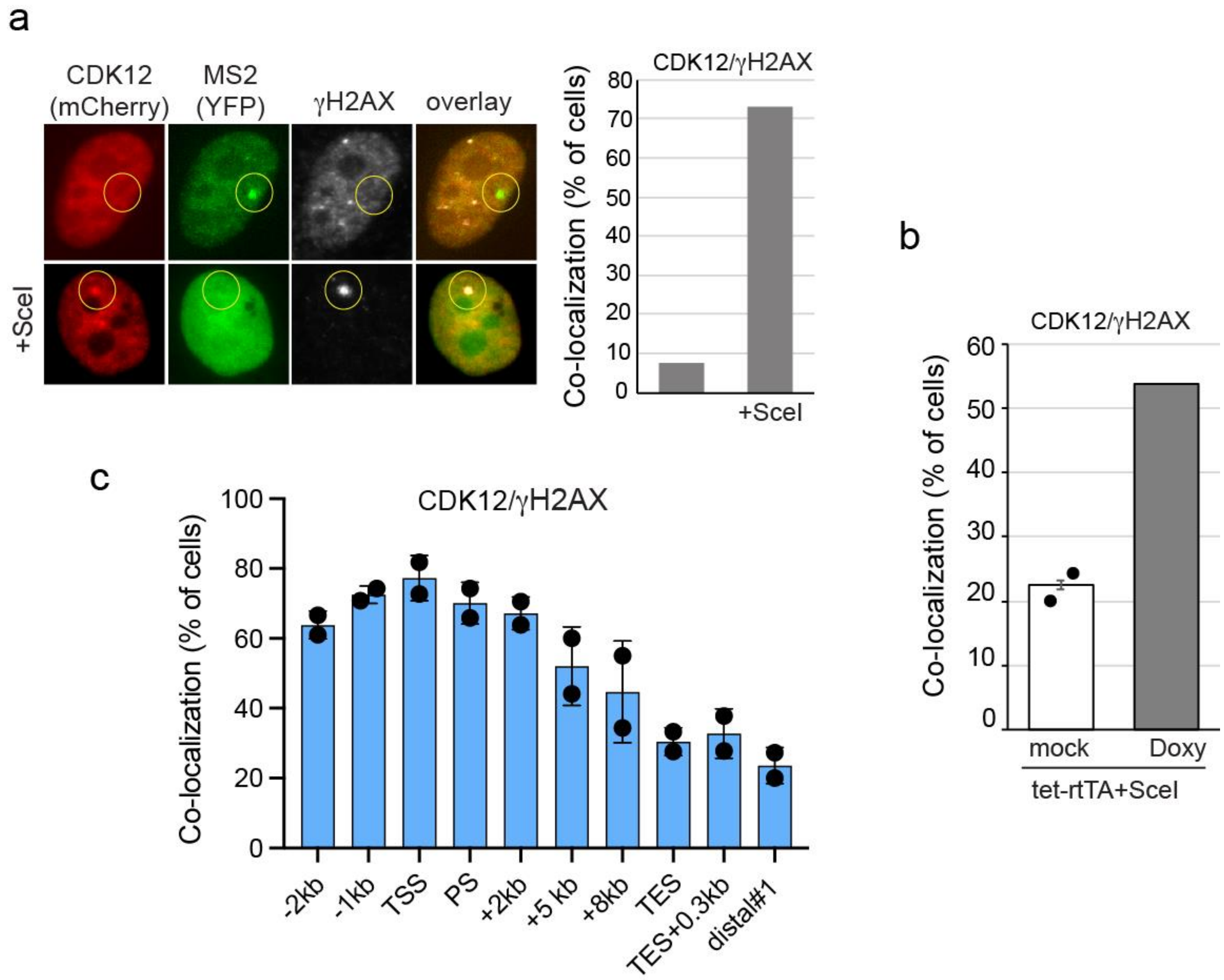


**Extended data figure 19. CDK12 colocalizes with DSBs at transcribed genes.**

**a**, CDK12 is recruited to the TRE-MS2 reporter only when DDR foci (γH2AX) are induced (SceI+). Left, snapshots U2OS-TRE-I-SceI-19 cells transfected with mCherry-CDK12 and YFP-MS2. Right, bar plot. (n=40; +SceI n=70).

**b**, Colocalization of γH2AX and CDK12 at the TRE-MS2 reporter in cells co-transfected with tet-rtTA and I-SceI. rtTA expression was induced with doxycycline (Doxy) for 5 hours. Mock: n=49, n=195; Doxy: n=230.

**c**, Colocalization of γH2AX and CDK12 along the MCM2 gene at sites where DSBs were introduced by CRISPR-Cas9 editing. TSS: transcription start site. TES: transcription termination site. Distal1: gene desert, non-transcribed genomic locus. Average of two independent experiments. -2kb: n=23, n=27; -1kb: n=24, n=35; TSS: n=44, n=44; PS: n=41, n=35; +2kb: n=36, n=44; +5kb: n=34, n=30; +8kb: n=32, n=40; TES: n=29, n=18; TES+0.3kb: n=36, n=37; distal#1: n=19, n=30.

*Extended data figure 20*

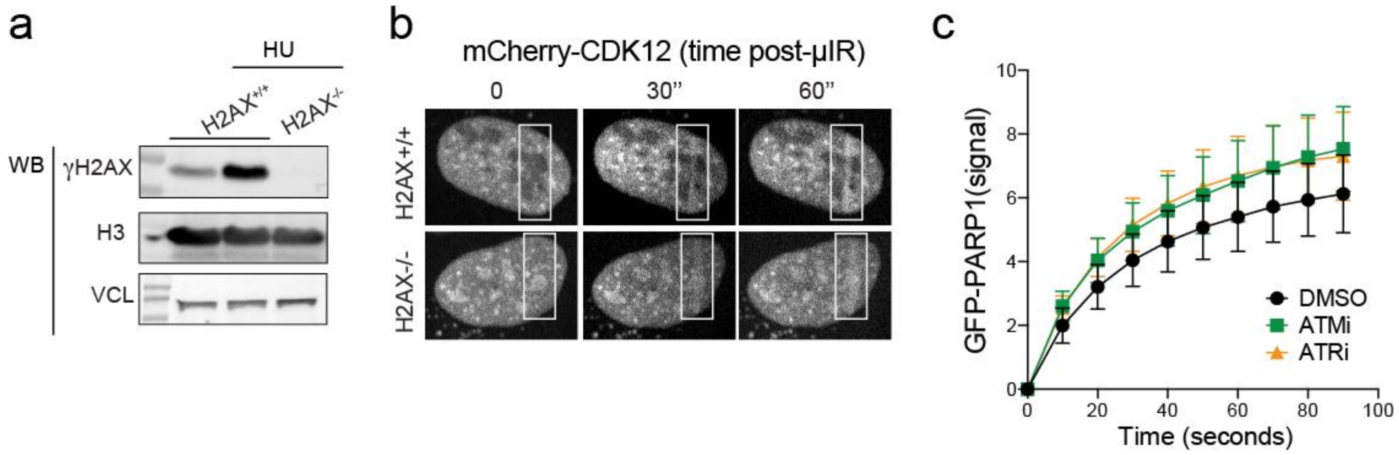


**Extended data figure 20. Recruitment of CDK12 is regulated by PARP.**
**a**, WB analysis of wild-type and H2AX knock-out U2OS cells.
**b**, Representative images showing the recruitment of mCherry-CDK12 to laser-damaged sites in H2AX knock-out cells.
**c**, Kinetics of PARP1 recruitment at laser-irradiated nuclear areas in cell treated with either ATMi (10 µM KU-5593) or ATRi (1 µM VE-821). DMSO: n=19; ATMi: n=18; ATRi: n=9.

*Extended data figure 21*

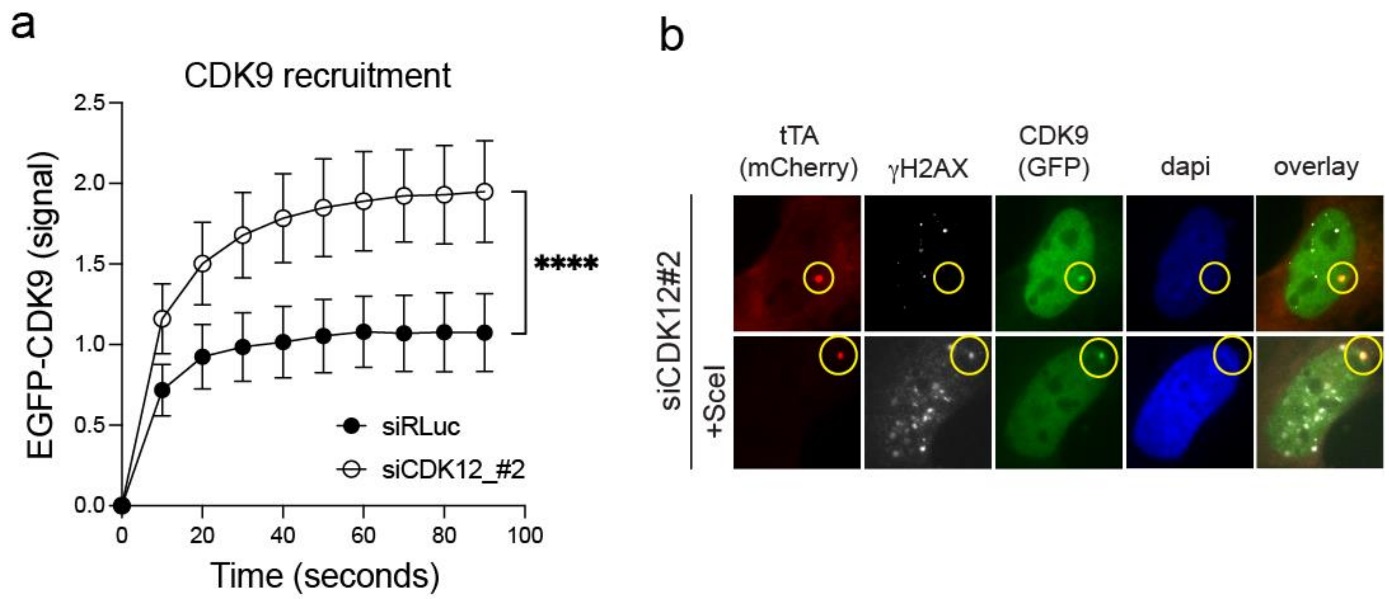


**Extended data figure 21. Silencing of CDK12 enhances the recruitment of CDK9 to damaged DNA.**
**a**, Kinetics of the recruitment of GFP-CDK9 at laser-damaged DNA. siRLuc: n=23; siCDK12#2: n=18.
**b**, Representative pictures showing that loss of CDK12 expression triggers the colocalization of GFP-CDK9 on the DNA damaged (+SceI) reporter locus of U2OS-TRE-I-SceI-19 cells.

*Extended data figure 22*

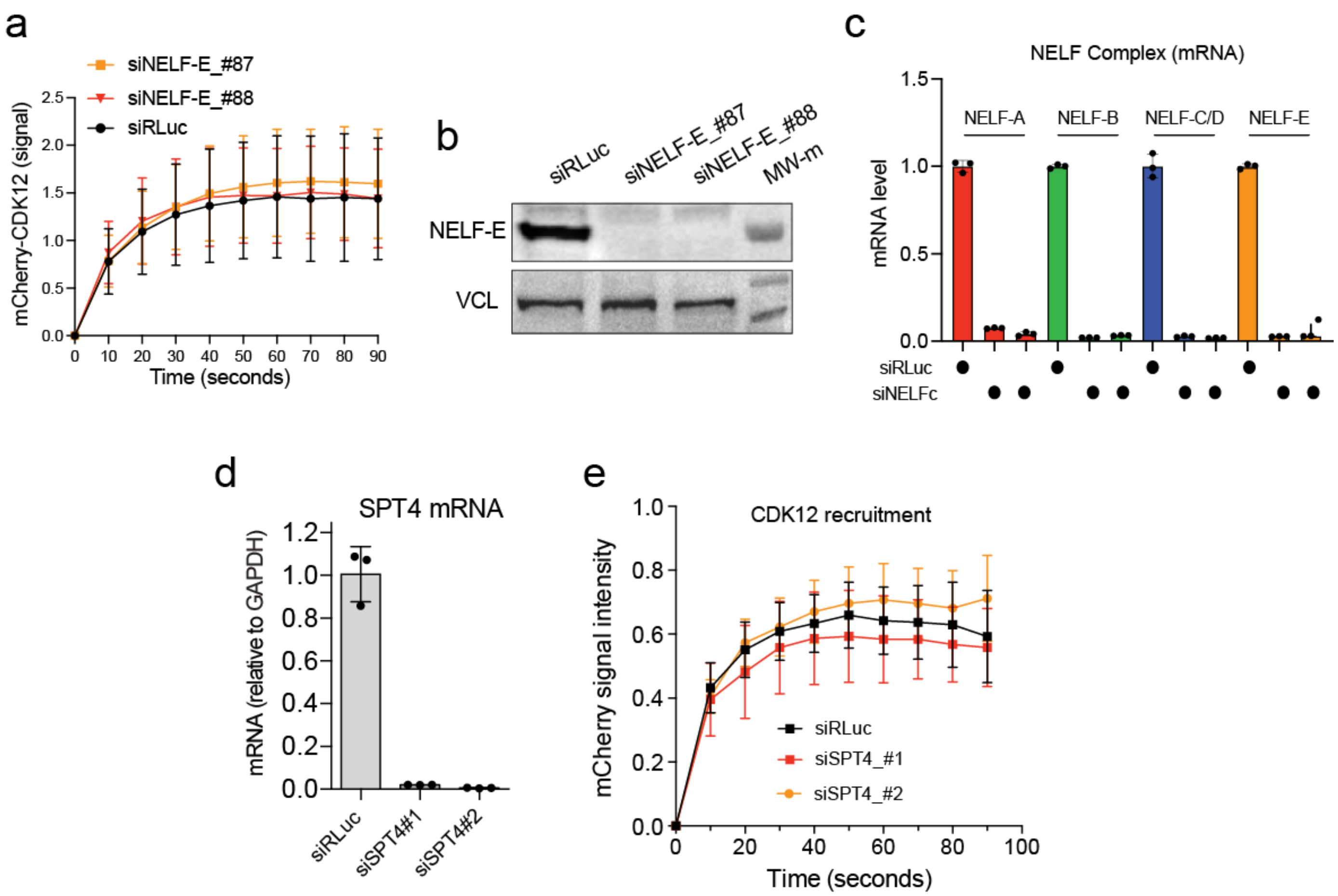


**Extended data figure 22. NELF complex does not regulate CDK12 recruitment to DNA-damaged sites.**
**a**, Silencing the NELF-E subunit does not affect mCherry-CDK12 recruitment to laser-damaged DNA. siRLuc: n=15; siNELF-E_#87: n=23; siNELF-E_#88: n=9.
**b**, WB analysis of U2OS cells transfected with siRNAs targeting NELF-E.
**c**, NELF-A, NELF-B, NELF-C and NELF-E mRNA expression evaluated by RT-qPCR following their silencing.
**d**, Bar plot of the average mRNA expression measured by RT-qPCR (n=3, error bar=stdv.).
**e**, Recruitment of mCherry-CDK12 to laser-damaged DNA in U2OS cells transfected with siRNAs targeting SPT4. siRLuc: n=16; siSPT4#1: n=17; siSPT4#2: n=17.

*Extended data figure 23*

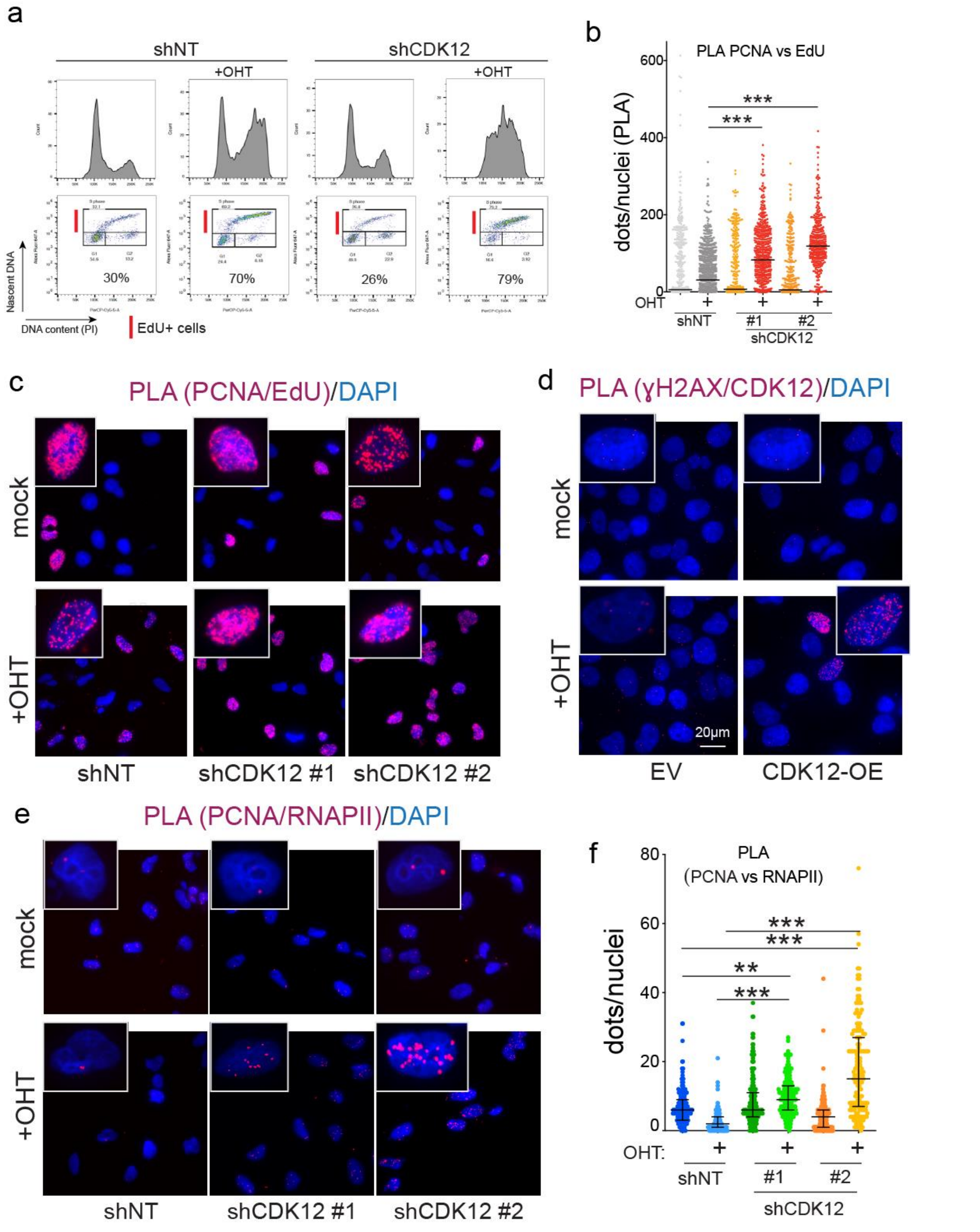


**Extended data figure 23. Silencing of CDK12 triggers replicative stress and TRCs upon MycER activation**.

**a**, Cell cycle entry of U2OS-MycER cells analyzed by FACS. Cells were pulse labeled with EdU and collected at 18 hours post-mitotic release. **b**, Bee swarm plot of PLA for PCNA and nascent DNA (EdU labeled). shNT: n=464 (mock), n=703 (+OHT), shCdk12#1: n=516 (mock), n=539 (+OHT), shCDK12#2: n=513 (mock), n=362 (+OHT). **c**, Images of PLA for PCNA and nascent DNA (EdU labeled). **d**, Representative images of PLA of CDK12 and yH2AX in U2OS-MycER cells. Where indicated (Cdk12-OE), cells were transfected with a plasmid encoding CDK12. **e**,**f**, PLA of PCNA and RNAPII in U2OS-MycER cells. shNT: n=214 (mock), n=136 (+OHT), shCdk12#1: n=186 (mock), n=194 (+OHT), shCDK12#2: n=226 (mock), n=167 (+OHT).

*Extended data figure 24*

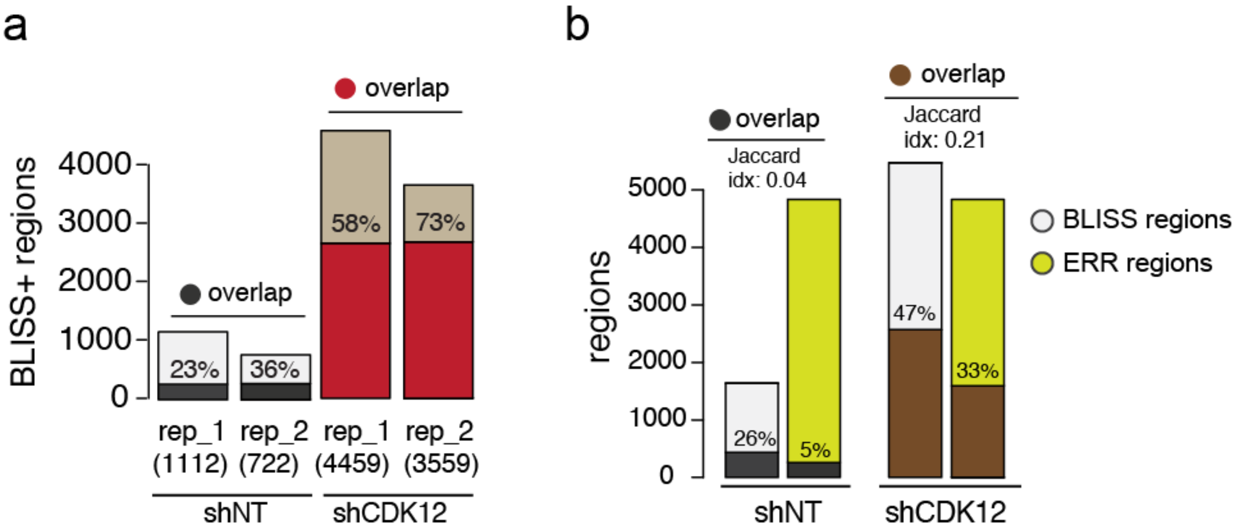


**Extended data figure 24. Genome-wide mapping of DSB-clusters (BLISS+ regions) upon MYC activation.**

**a**, BLISS+ regions identified in replicate experiments in U2OS-MycER cells upon MycER activation and CDK12 silencing.

**b**, Bar plot showing the overlap of the ERR+ and the BLISS+ regions in U2OS-MycER cells upon MycER activation and CDK12 silencing.

*Extended data figure 25*

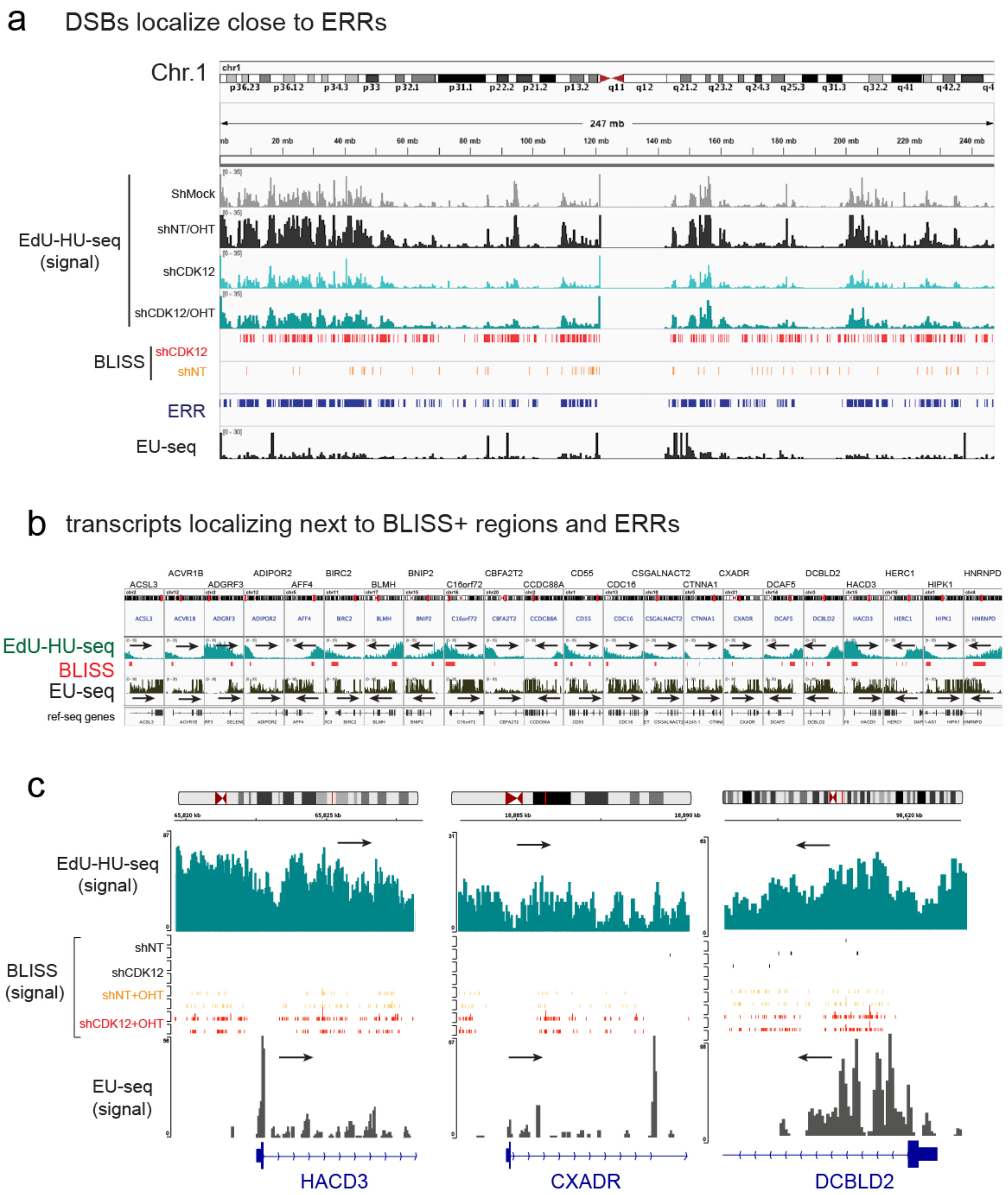


**Extended data figure 25. Genomic view of BLISS+ regions.**

**a**, Genome browser view of chromosome 1 showing how DSBs (BLISS+ regions) localize in proximity of early replicated regions (ERRs) mapped by EdU-HU-seq. **b**, Genome browser snapshots of genes at ERRs showing EdU-HU-seq (DNA replication) and EU-seq (Nascent RNA) signals along with the position of the BLISS+ regions. Black arrows indicate direction of DNA and RNA synthesis. This shows that BLISS+ regions are nested between codirectional ERRs and the promoter of transcribed genes. **c**, Genomic snapshots of genes reporting the Bliss signals (reads) detected in U2OS-MycER cells upon mock silencing (shNT), MycER activation (shNT + OHT), CDK12 silencing (shCDK12) and CDK12 silencing and MycER acrtivation (shCDK12 + OHT).

*Extended data figure 26*

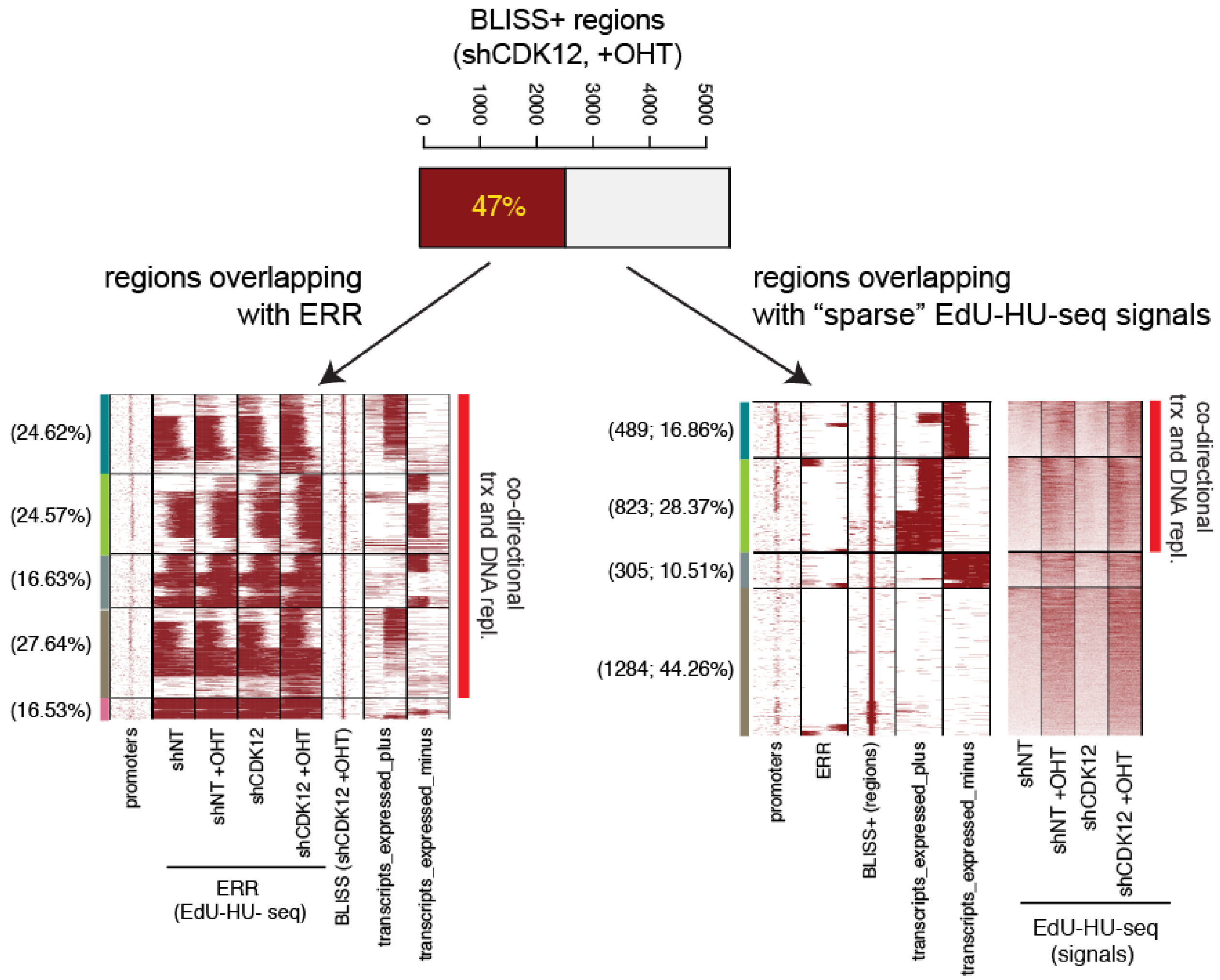


**Extended data figure 26. Workflow for the classification of BLISS+ region identified in shCDK12 MycER cells.** BLISS+ regions were first subsetted based on their overlap with ERR+ (47%) and then were clustered based on the ERR+ and presence of transcripts. The same strategy was used to cluster the ERR- regions. Analysis of the EdU-HU-seq data revealed that these latter regions have positive EdU-HU-seq signals deriving from sparse DNA replication. Hence these regions were named “overlapping with sparse EdU-HU signals”.

*Extended data figure 27*

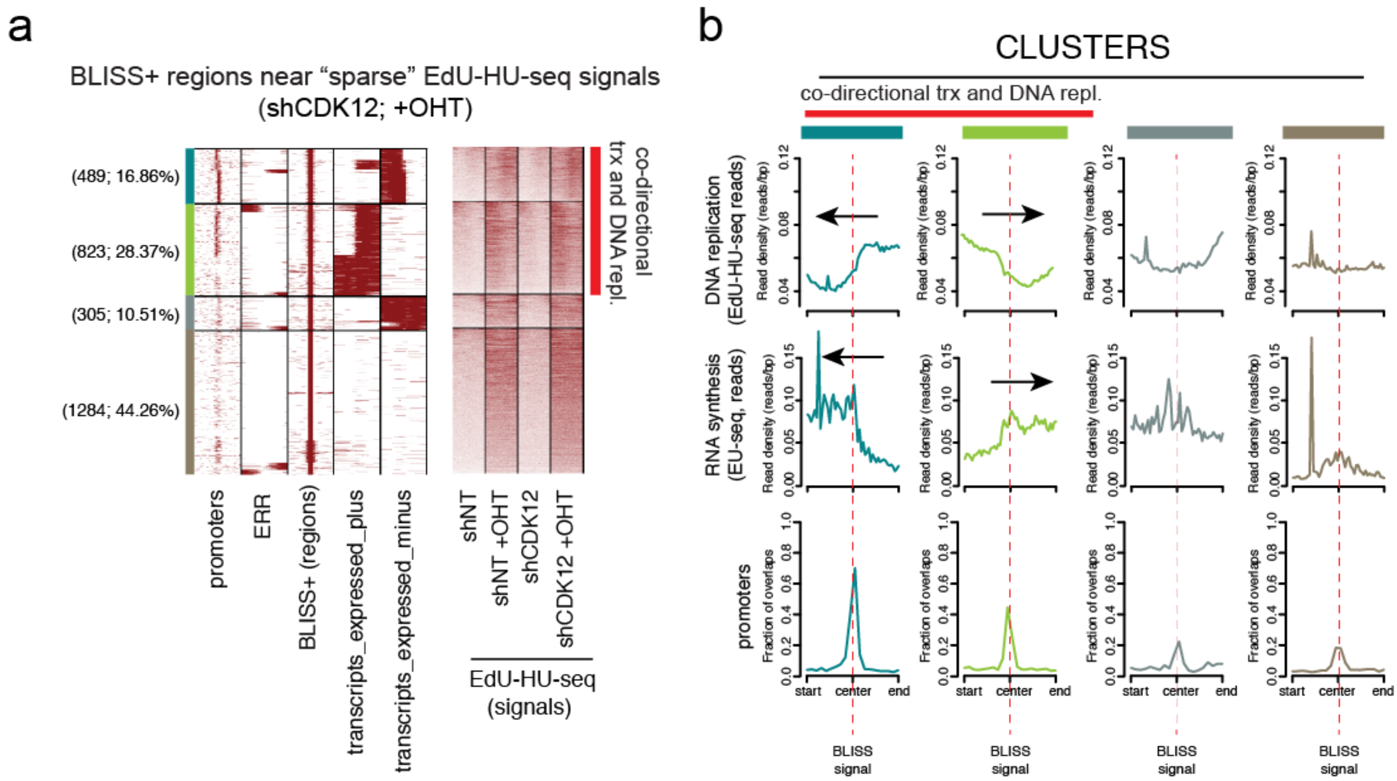


**Extended data figure 27. BLISS+ regions neither proximal nor overlapping with ERRs identified in U2OS cells upon MycER activation and CDK12 silencing.**

**a**, Clustered heatmap of genomic regions (10kb) centered on the BLISS signal.

**b**, EdU-HU-seq and EU-seq signal distribution plot and promoter frequency distribution in the different cluster identified in (a). Arrows indicate the direction of DNA and RNA synthesis in the two clusters with co-directional transcription and DNA replication, which account for 45% of the BLISS+ regions near sparse EdU-HU-seq signals.

*Extended data figure 28*

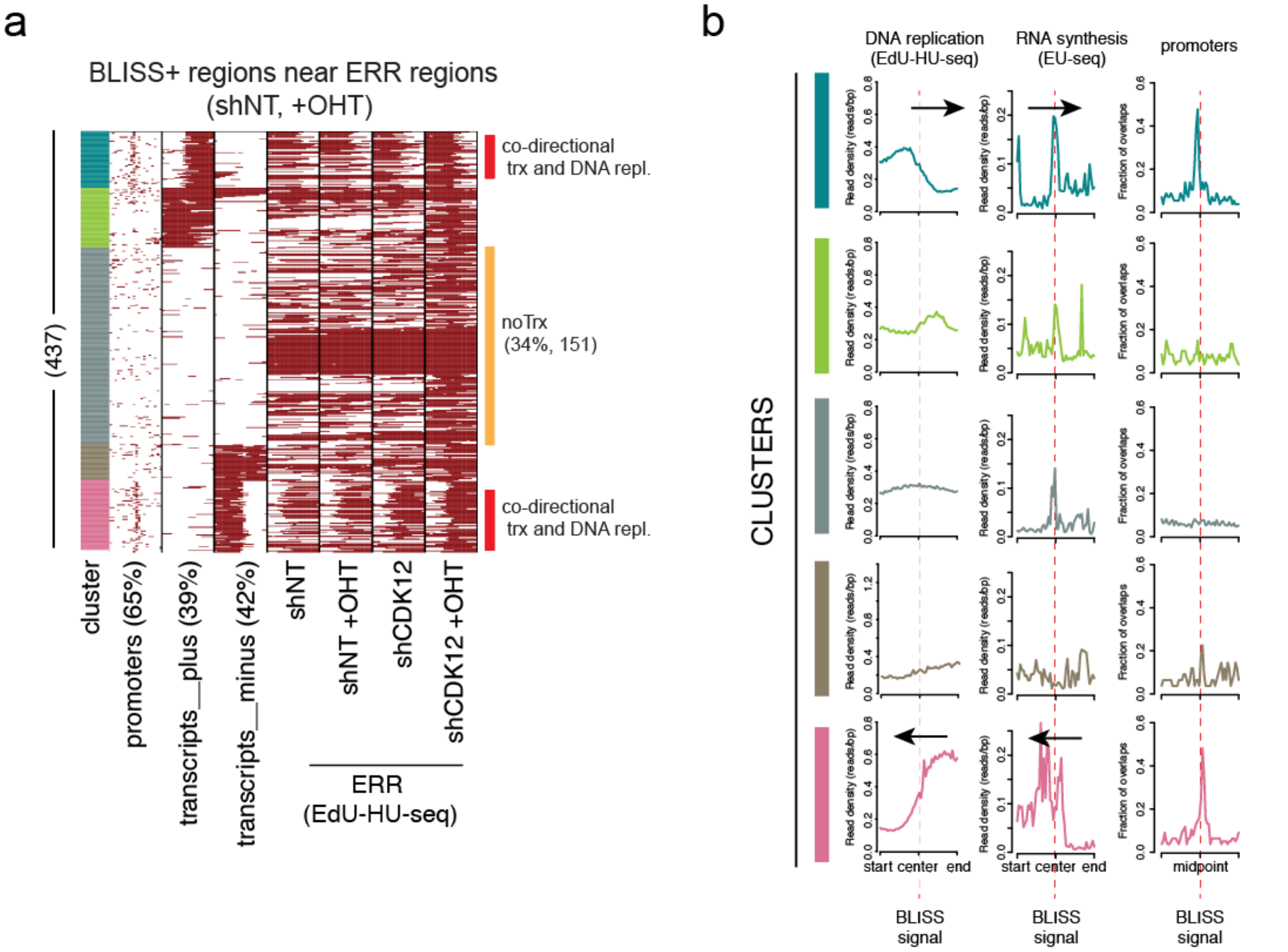


**Extended data figure 28. BLISS+ regions proximal or overlapping with ERRs identified in U2OS cells upon MycER activation.**
**a**, Clustered heatmap of genomic regions (10kb) centered on the BLISS+ regions
**b**, EdU-HU-seq and EU-seq signal distribution plot and promoter frequency distribution in the different cluster identified in (a). Arrows indicate the direction of DNA and RNA synthesis in the two clusters with codirectional transcription and DNA replication, which account for 30% of the BLISS+ regions near ERR.

### Extended data figure 29

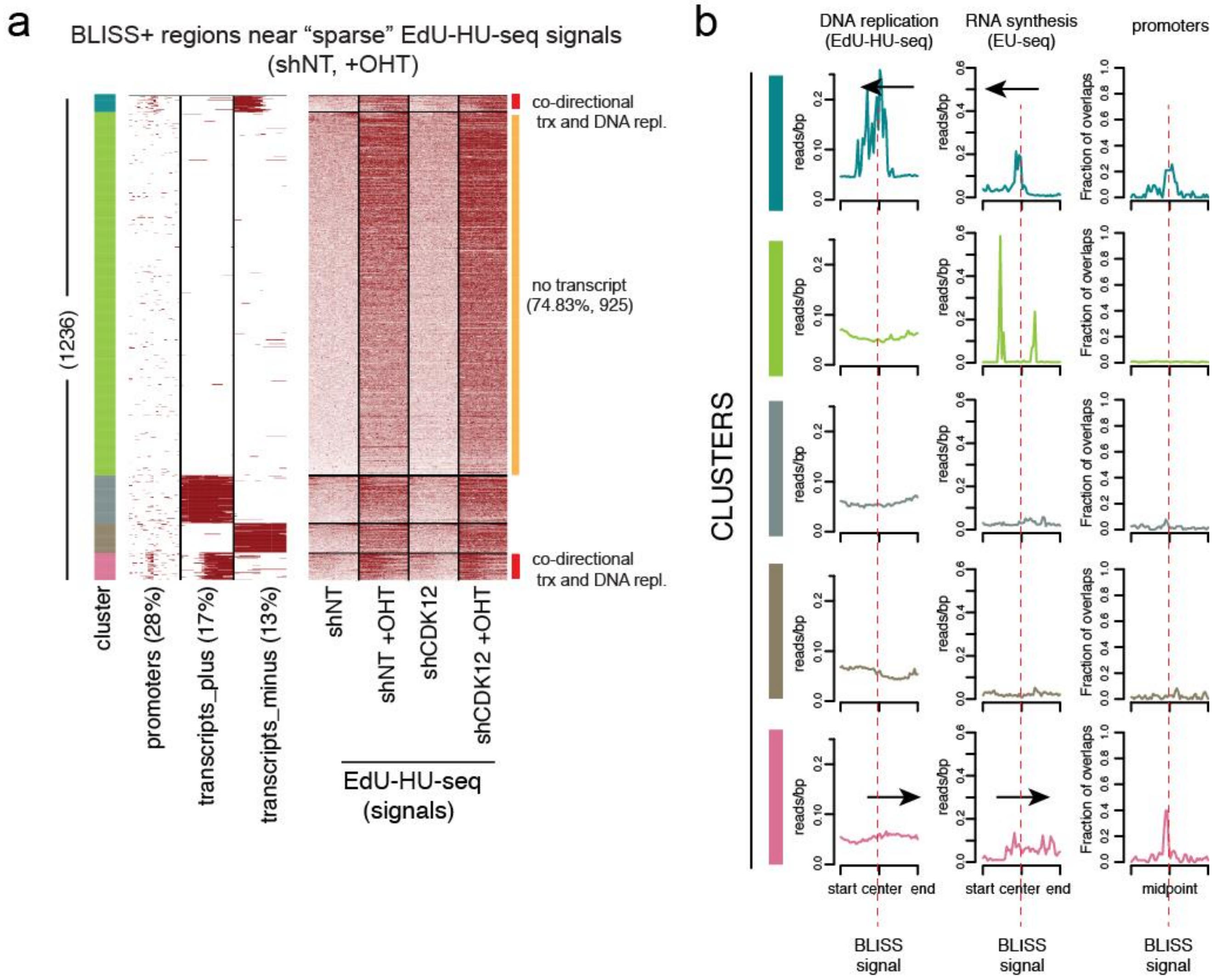


**Extended data figure 29. BLISS+ regions neither proximal nor overlapping with ERRs identified in U2OS cells upon MycER activation in mock silenced cells.**
**a**, Clustered heatmap of genomic regions (10kb) centered on the BLISS+ region.
**b**, EdU-HU-seq and EU-seq signal distribution plot and promoter frequency distribution in the different cluster identified in (**a**). Arrows indicate the direction of DNA and RNA synthesis in the two clusters with codirectional transcription and DNA replication, which account for 9% of the BLISS+ regions near ERR.

*Extended data figure 30*

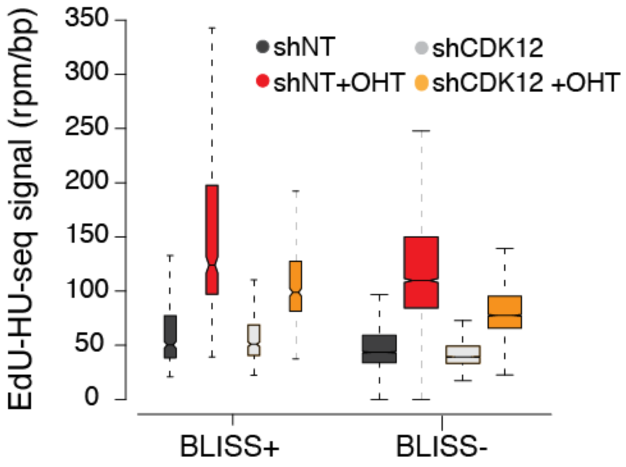


**Extended data figure 30. MYC stimulates stronger DNA replication on ERRs prone to undergo DSBs.** Box plot of EdU-HU-seq signals on the left side of ERRs overlapping with BLISS+ regions (BLISS+) or not (BLISS-).

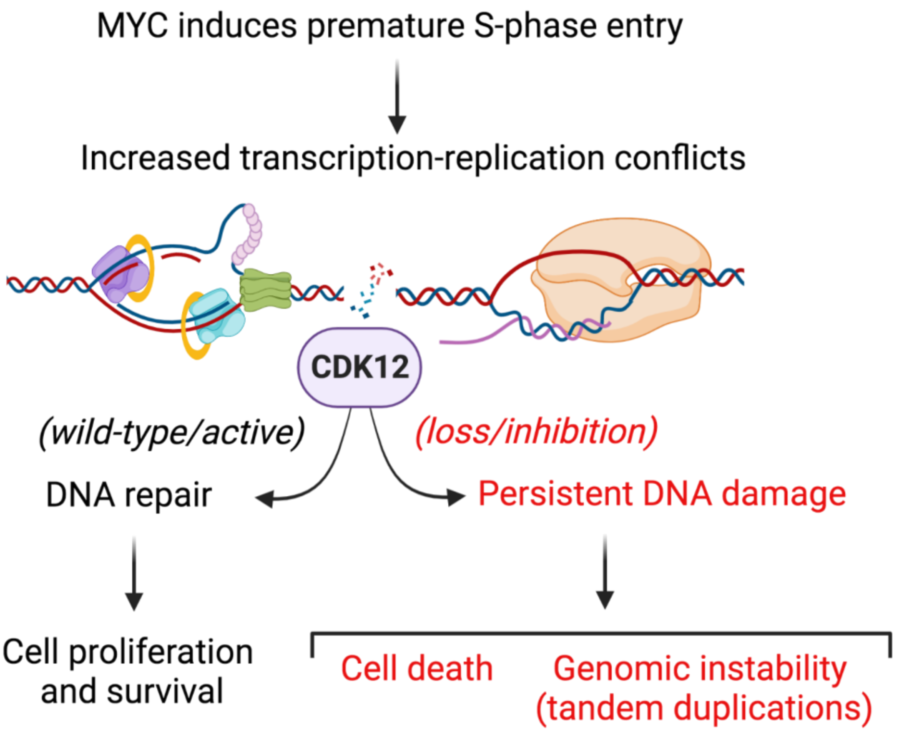


**Extended data figure 31. Model of how CDK12 may prevent genomic instability by favoring resolution of DSBs at TRCs.** Created with BioRender.com.

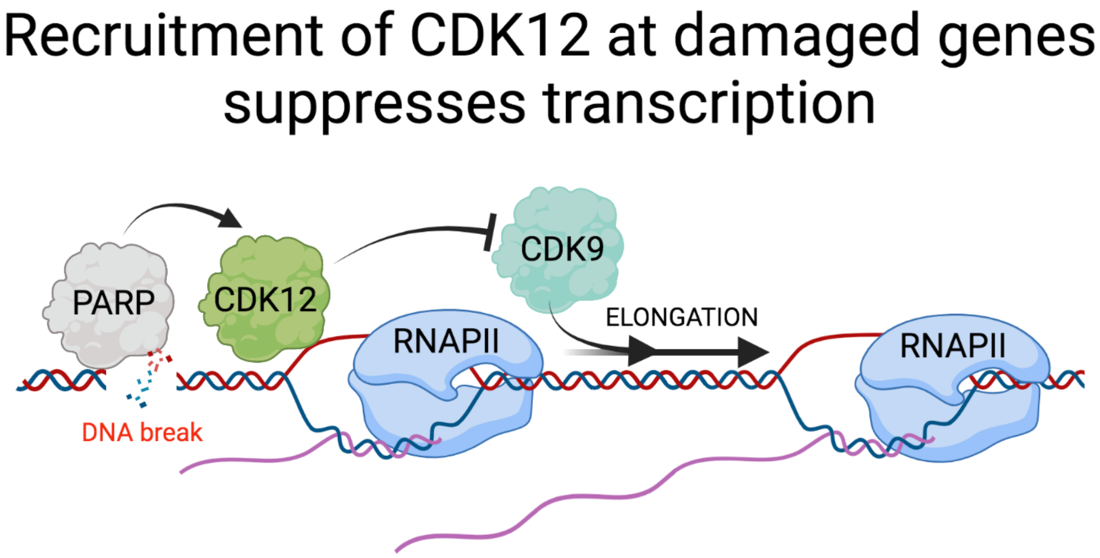


**Extended data figure 32. Scheme of how CDK12 is recruited at damaged genes to regulate their transcription.** Created with BioRender.com.
